# The native cell differentiation program aberrantly recapitulated in *yki*^*3S*/*A*^-induced intestinal hyperplasia drives invasiveness and cachexia-like wasting phenotypes

**DOI:** 10.1101/2022.06.01.494390

**Authors:** Inez K.A. Pranoto, Jiae Lee, Young V. Kwon

## Abstract

Many tumors recapitulate the developmental and differentiation program of their tissue of origin, a basis for tumor cell heterogeneity. Although stem-cell-like tumor cells are well-studied, the roles of tumor cells undergoing differentiation in inducing the phenotypes associated with advanced cancers remains to be elucidated. Here, we employ *Drosophila* genetics to demonstrate that the native differentiation program of intestinal stem cells plays a key role in determining an intestinal tumor’s capacity to invade and induce various non-tumor-autonomous phenotypes. The differentiation program that generates absorptive cells enterocytes is aberrantly recapitulated in the intestinal tumors generated through activation of the Yap1 ortholog Yorkie. Elimination of tumor cells in the enterocyte lineage allows stem cell-like tumor cells to grow but suppresses invasiveness and reshapes various phenotypes associated with cachexia-like wasting by altering the expression of tumor-derived factors. Our study provides insight into how a native differentiation program determines a tumor’s capacity to induce the phenotypes associated with advanced cancers and suggests that manipulating the differentiation programs co-opted in tumors might be a way to treat some complications of cancer, including cachexia.

## Introduction

Division of stem cells and their differentiation processes generate the heterogeneous cell populations required for tissue development and maintenance (Hwang et al., 2008; Jiang et al., 2009; Jiang et al., 2016; Krieger and Simons, 2015; Micchelli and Perrimon, 2006; Ohlstein and Spradling, 2006; Zakrzewski et al., 2019). Recent studies employing single-cell sequencing strategies have highlighted the striking cellular heterogeneity of human cancers and mouse cancer models (Couturier et al., 2020; Goveia et al., 2020; Kim et al., 2020; Lawson et al., 2018; Wu et al., 2021; Yeo et al., 2020). In particular, several human cancers have been shown to recapitulate the developmental and/or differentiation programs that form and maintain the tissues of origin (Borczuk et al., 2003; Couturier et al., 2020; Fukuzawa et al., 2017; Goveia et al., 2020; Kelleher et al., 2006; Wu et al., 2021). Interestingly, Genovese et al. have shown that the temporal patterning program is partially co-opted in *Drosophila* neuroblast tumors (Genovese et al., 2019). Along with the concept that dedifferentiation is associated with malignancy, the roles of cancer-cell populations with stem cell-like properties have been extensively studied (Chen et al., 2013; da Silva-Diz et al., 2018; Jögi et al., 2012). It remains to be determined, however, how these developmental and differentiation programs recapitulated in cancers contribute to the various phenotypes associated with advanced cancers, including metastasis and cachexia, most of which are responsible for the mortality of cancer patients.

Oncogenes do not elicit the same tumorigenicity and tumor-related phenotypes across different tissues (Cook et al., 2021; Figueroa-Clarevega and Bilder, 2015; Kwon et al., 2015; Lee et al., 2020; Lowell et al., 2000; Pagliarini and Xu, 1991; Rangarajan et al., 2001; Tu et al., 2019; Weng et al., 2006; Yang et al., 2019), implying that the characteristics of cancers cannot be attributed entirely to the alterations in their genome. Similarly, in *Drosophila*, expression of an oncogene often give rise to different phenotypes in the imaginal discs–the developing epithelia inside larvae–and the adult midgut epithelium. Expression of activated forms of the Yes-associated protein 1 (Yap1) oncogene ortholog *yorkie* (*yki*), such as *yki*^*S168A*^ and *yki*^*3S/A*^, is sufficient to drive uncontrolled cell division in multiple epithelial tissues, resulting in the formation of hyperplasia (hereafter, *Drosophila* ‘tumors’) (Kwon et al., 2015; Oh and Irvine, 2009; Pan, 2010; Wang et al., 2016; Wittkorn et al., 2015). Transplantation of *yki*^*S168A*^ imaginal disc tumors into adult flies does not induce wasting in the host tissues even though the transplanted tumors grow large inside the hosts (Figueroa-Clarevega and Bilder, 2015). In contrast, formation of *yki*^*3S/A*^ tumors in the midgut epithelium induces cachexia-like wasting, manifested by ovary atrophy, muscle degeneration, and metabolic abnormalities (Kwon et al., 2015). Additionally, while imaginal disc tumors generated by activation of Yki are not metastatic, our observations show that a portion of *yki*^*3S/A*^ midgut tumor cells can migrate out of the midgut across the visceral muscle (VM). These observations suggest that the malignant phenotypes caused by *yki*^*3S/A*^ midgut tumors cannot be attributed solely to the activation of Yki. It is plausible to speculate that midgut-specific contextual information might play a role in eliciting these phenotypes. In this study, we attempted to address how the invasive and the cachexia-like wasting phenotypes arise in *yki*^*3S/A*^ midgut tumors in order to gain insights into how the physiology of the tissue of origin contributes to a tumor’s ability to induce malignant phenotypes.

While the epithelial cells in imaginal discs are relatively homogenous, the intestinal epithelium comprises 4 cell types: intestinal stem cells (ISCs), enteroblasts (EBs), enterocytes (ECs), and enteroendocrine cells (EEs) (Boumard and Bardin, 2021; Hou and Singh, 2017; Jiang et al., 2016; Micchelli and Perrimon, 2006; Miguel-Aliaga et al., 2018; Ohlstein and Spradling, 2006). ISCs are the diploid cells expressing Delta (Dl)–the ligand of the Notch (N) signaling pathway. ISCs divide and by default generate themselves while activation of Notch signaling makes ISCs generate their progenitor cells EBs (Boumard and Bardin, 2021; Hou and Singh, 2017; Jiang et al., 2016; Micchelli and Perrimon, 2006; Miguel-Aliaga et al., 2018; Ohlstein and Spradling, 2006). Subsequent activation of Notch signaling in EBs triggers differentiation of EBs into ECs, which are the absorptive polyploid cells (Micchelli and Perrimon, 2006). ISCs also undergo a distinct lineage to give rise to EEs (Biteau and Jasper, 2014; Zeng and Hou, 2015). The midguts generate intermediately-differentiated EBs, which then migrate and terminally differentiate to replace damaged ECs, a process requiring epithelial-mesenchymal transition (EMT) followed by mesenchymal-epithelial transition (MET). A snail family EMT transcription factor, *escargot* (*esg*), plays a key role in EC differentiation (Antonello et al., 2015; Korzelius et al., 2014; Loza-Coll et al., 2014). ISCs express less *esg* compared to EBs, which makes ISCs be at a partial EMT state while EBs elicit a full EMT (Antonello et al., 2015). Indeed, the differentiating EBs show the morphological features of mesenchymal cells and the ability to migrate (Antonello et al., 2015). Note that stem cells can also migrate upon tissue damage (Hu et al., 2021). Although midgut tumors can be generated by expressing oncogenes in ISCs and EBs (Apidianakis et al., 2009; Lee et al., 2021; Markstein et al., 2014), it is not clear whether the normal differentiation programs are recapitulated in these tumors to generate a heterogeneous population of tumor cells. Furthermore, it remains underexplored how a differentiation program recapitulated in tumors or how a population of tumor cells undergoing differentiation contributes to the expression of the various advanced tumor phenotypes in *Drosophila* as well as in humans.

We started our study by describing a subpopulation of *yki*^*3S/A*^-induced midgut tumor cells (*yki*^*3S/A*^ cells) that showed invasive phenotypes, leading us to hypothesize that *yki*^*3S/A*^ cells were not homogeneous. Our characterization revealed the striking heterogeneity of *yki*^*3S/A*^ cells, which aberrantly reconstituted the EC differentiation program. By genetically blocking EC differentiation in *yki*^*3S/A*^ tumors, we demonstrated that maintaining the EC differentiation program was crucial for the ability of *yki*^*3S/A*^ cells to disseminate from the midguts and induce certain non-tumor-autonomous phenotypes. Our study provides insights into how the contextual information in the tissue of origin can give rise to a tumor’s capacity to induce the phenotypes associated with invasiveness and cachexia-like wasting.

## Results

### A portion of *yki*^*3S/A*^ cells basally disseminate from the posterior midguts

We previously showed that expression of a mutant *Ras* (*Ras*^*V12*^) in adult ISCs and EBs using the conditional GAL4 driver *esg*^*ts*^ (*esg-GAL4, tub-GAL80*^*ts*^, *UAS-GFP*/+; see Methods) could induce tumors in the midguts for a short period but at day 2 of *Ras*^*V12*^ expression, they basally disseminate and apically delaminate, resulting in a removal of most of *Ras*^*V12*^ cells from the posterior midguts (Lee et al., 2020). In contrast, expression of *yki*^*3S/A*^ with *esg*^*ts*^ (*esg*^*ts*^*>yki*^*3S/A*^) resulted in midgut tumors which persist over time (Fig. 1A and S1A) (Kwon et al., 2015). Unlike *Ras*^*V12*^ cells at day 2, *yki*^*3S/A*^ cells showed strong Armadillo (Arm)–the *Drosophila* β-catenin ortholog–signals at the cell-cell junctions, an indication of intact adherens junctions (Fig. 1A). Moreover, overall Arm signals were significantly increased in *yki*^*3S/A*^ cells compared to control *esg*^*+*^ cells (ISCs and EBs) (Fig. 1A and B). Consistently, most of the *yki*^*3S/A*^ cells stayed at the epithelium, and apical delamination of *yki*^*3S/A*^ cells was not frequently observed (Fig. S1B). Strikingly, a significant number of *yki*^*3S/A*^ cells were also detected outside of the VM at the posterior midguts as early as day 4 of *yki*^*3S/A*^ expression, a phenotype that became more robust over time (Fig. 1C and D). These observations indicate that a portion of *yki*^*3S/A*^ cells can invade and migrate out from the midguts across the VM and the extracellular matrix (ECM) while most of *yki*^*3S/A*^ cells stay in the epithelium.

**Figure 1.**
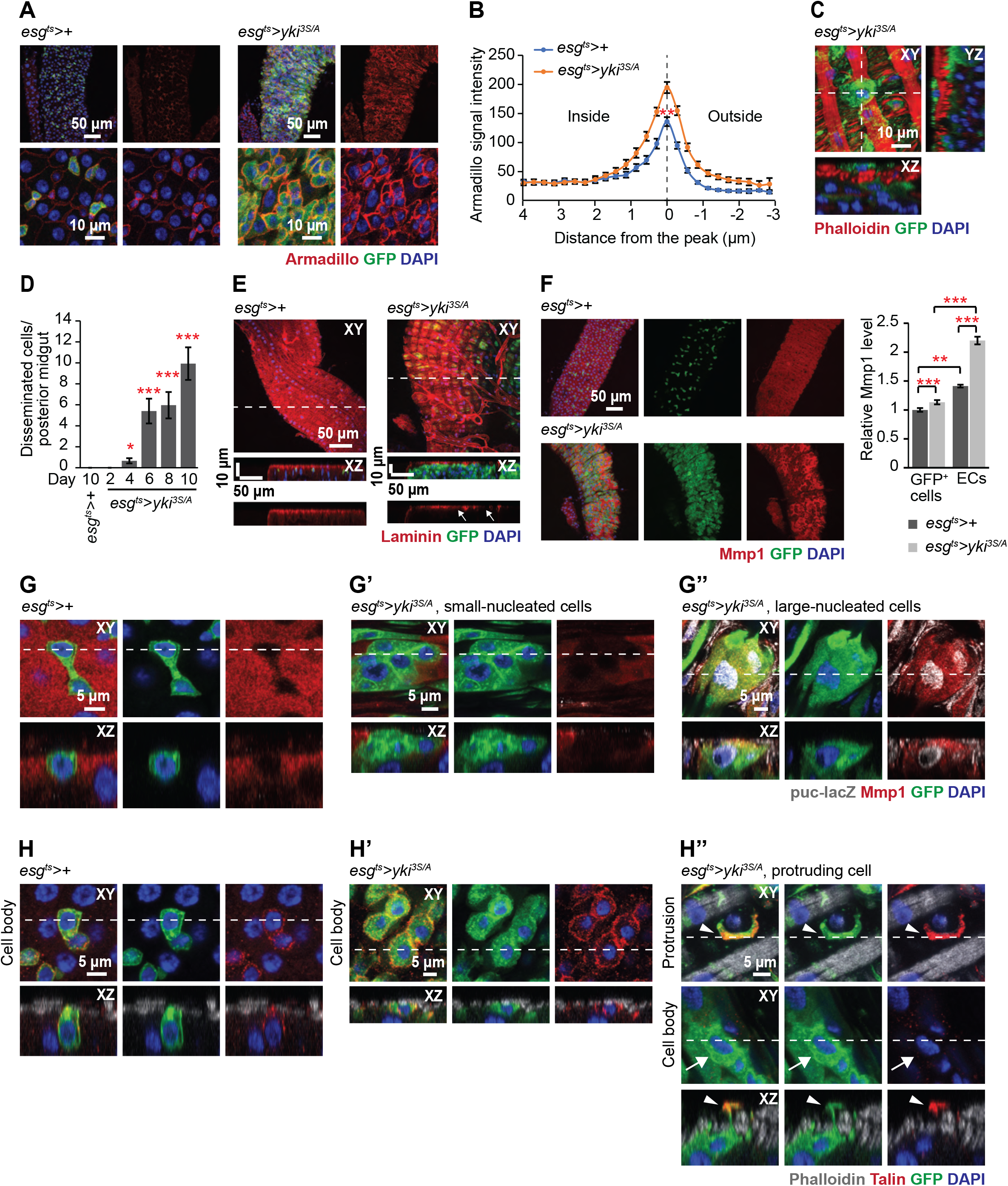
A portion of cells in *yki*^*3S/A*^-induce midgut hyperplasia disseminate from the midguts. **A**. Arm signals (red) in the posterior midguts. Transgenes were induced 6 days using *esg-GAL4, UAS-GFP, Tub-GAL80*^*ts*^ (*esg*^*ts*^) by shiting to 29°C. Cells manipulated by *esg*^*ts*^ are marked by GFP (green). Nuclei were stained by DAPI (blue). **B**. Quantification of Arm signals in *GFP*^*+*^ cells. Transgenes were induced for 6 days. N=31 cells from 5 midguts, *esg*^*ts*^*>+*; N=39 cells from 11 midguts, *esg*^*ts*^>*yki*^*3S/A*^. **P<0.01, chi-square test. **C**. Representative image of disseminated *yki*^*3S/A*^ cell. Top and orthogonal views are shown. Phalloidin (red) visualizes visceral muscle. **D**. Quantification of disseminated cells. *GFP*^*+*^ and *DAPI*^*+*^ cells detected outside the VM layer in the posterior midguts are counted. Transgenes were induced for indicated days with *esg*^*ts*^. N=22 midguts (10 days, *esg*^*ts*^*>+*); N=24 (2 days, *esg*^*ts*^>*yki*^*3S/A*^); N=24 (4 days, *esg*^*ts*^>*yki*^*3S/A*^); N=32 (6 days, *esg*^*ts*^>*yki*^*3S/A*^); N=28 (8 days, *esg*^*ts*^>*yki*^*3S/A*^); N=29 (10 days, *esg*^*ts*^>*yki*^*3S/A*^). Mean±SEMs are shown, and *P<0.05, **P<0.01, and ***P<0.001 by two-tailed unpaired Student’s t-test. **E**. Laminin staining (red) of the midguts. Arrows indicate localized degradation of the laminin layer. **F**. Mmp1 staining (red) at 6 days of transgene induction. The graph shows quantification of Mmp1 signals in *GFP*^*+*^ and EC cells. N=90 cells from 9 *esg*^*ts*^*>+* midguts; N=110 cells from 11 *esg*^*ts*^>*yki*^*3S/A*^ midguts. Mean±SEMs are shown, and *P<0.05, **P<0.01, and ***P<0.001 by two-tailed unpaired Student’s t-test. **G-G’’**. Mmp1 and *puc-lacZ* signals in the midgut epithelia. Representative images of control cells (*esg*^*ts*^*>+*) (**G**), *yki*^*3S/A*^ cells without Mmp1 and *puc-lacZ* signals (**G’)** and *yki*^*3S/A*^ cells with Mmp1 and *puc-lacZ* signals (**G’’**) are shown. In the last panels, both Mmp1 (red) and *puc-lacZ* signals (gray) are shown. **H-H’’**. Talin localization in *esg*^*+*^ cell in control (*esg*^*ts*^*>+*) and *esg*^*ts*^>*yki*^*3S/A*^ midguts. Representative images of control cells (**H**), *yki*^*3S/A*^ cells without forming protrusions (**H’**), and *yki*^*3S/A*^ cell forming protrusions (**H’’**) are shown. Phalloidin (gray) marks the visceral muscle. “Cell body panel” shows a z-stack spanning nuclei of *GFP*^*+*^ cells. “Protrusion panel” shows a view from outside. Arrowhead indicates a protrusion and arrow shows the corresponding cell body.

### Matrix metalloproteinase 1 (Mmp1) is increased only in a portion of *yki*^*3S/A*^ cells and most of ECs in *yki*^*3S/A*^ midguts

A relatively thick ECM layer exists at the basal side of the midgut epithelium (Howard et al., 2019; Lee et al., 2020). Thus, *yki*^*3S/A*^ cells need to breach the ECM to disseminate across the VM. To assess if the ECM was compromised by expression of *yki*^*3S/A*^ with *esg*^*ts*^, we stained control and *yki*^*3S/A*^ midguts with an anti-laminin antibody. In control midguts, one continuous laminin layer was detected at the basal side (Fig. 1E). In contrast, expression *yki*^*3S/A*^ with *esg*^*ts*^ resulted in a localized partial breach of the laminin layer (Fig. 1E, arrows). Next, we assessed Mmp1 levels in control and *yki*^*3S/A*^ midguts. In control midguts, Mmp1 signals were detected in ECs; *esg*^*+*^ cells did not show discernable signals (Fig. 1F and G). Interestingly, expression of *yki*^*3S/A*^ with *esg*^*ts*^ significantly increased Mmp1 signals in ECs in a non-cell-autonomous manner (Fig. 1F). Notably, only a portion of *yki*^*3S/A*^ cells showed elevated Mmp1 signals while Mmp1 signals were unaltered in most of the *yki*^*3S/A*^ cells (Fig. 1G’ and G’’). Thus, overall Mmp1 signals in *yki*^*3S/A*^ cells were increased slightly, yet significantly, when compared to those in control *esg*^*+*^ cells (Fig. 1F).

Prior studies have shown that c-Jun N-terminal Kinase (JNK) signaling increases Mmp1 in tumors and during wound healing in *Drosophila* (Ma et al., 2015; Stevens and Page-McCaw, 2012; Uhlirova and Bohmann, 2006). To address whether JNK signaling was associated with the elevation of Mmp1, we assessed JNK signaling using *puc-lacZ*, which expresses nuclear LacZ under the control of the regulatory sequence of *puckered* (*puc*)–a feedback regulator of JNK signaling (Martín-Blanco et al., 1998). We found that *yki*^*3S/A*^ cells with increased Mmp1 signals showed increased LacZ signals (Fig. 1G’’) while LacZ signals were rarely detected in control *esg*^*+*^ cells (Fig. 1G). Note that we attempted to address the role of Mmp1 in the dissemination of *yki*^*3S/A*^ cells by expressing an RNAi against Mmp1 (*JF01336*) or a dominant-negative Mmp1 (*Mmp1*^*DN Pro-pex*^) with *esg*^*ts*^ (Glasheen et al., 2009; Lee et al., 2012). Loss of Mmp1 or its activity decreased the growth of *yki*^*3S/A*^ tumors (Fig. S2A). Although Mmp1 depletion or inhibition completely suppressed dissemination of *yki*^*3S/A*^ cells, the defect in tumor growth likely accounted for the reduction (Fig. S2B).

These results show that expression of *yki*^*3S/A*^ with *esg*^*ts*^ partially degrades the laminin layer and increases Mmp1. Although expression of *yki*^*3S/A*^ with *esg*^*ts*^ significantly increases Mmp1 in most ECs, Mmp1 is elevated only in a portion of *yki*^*3S/A*^ cells. Given the role of Mmp1 in invasiveness, our observations indicate that only a portion of *yki*^*3S/A*^ cells is invasive.

### A portion of *yki*^*3S/A*^ cells form protrusions across the VM, which are enriched for focal adhesion components

In previous studies, we found that *Ras*^*V12*^ cells form bleb-like protrusions, which penetrate the VM by compromising the VM integrity (Lee et al., 2020). In contrast, here we found that only a portion of *yki*^*3S/A*^ cells formed elaborated protrusions while the majority did not (Fig. 1H-H’’). Additionally, these protrusions transversed the VM layer through the gaps between circular muscles without compromising the integrity of the tissue (Fig. S3A), suggesting that these protrusions are distinct from the bleb-like protrusions observed in *Ras*^*V12*^ cells.

During migration, cells often assemble focal adhesions at the leading edge, which serve as attachments for dragging the lagging cell bodies (Krakhmal et al., 2015; Pandya et al., 2017). To address whether focal adhesions were assembled at the protrusions transversing the VM, we first checked the localization of integrin–the transmembrane component of focal adhesions. In control cells (*esg*^*ts*^*>+*), Myospheroid (Mys)–a β-subunit of *Drosophila* integrin–signals were detected at the basal and the lateral sides of the cells (Fig. S3B). Similarly, the majority of *yki*^*3S/A*^ cells that were not forming protrusions showed Mys signals at the basal and the lateral sides with signals that were stronger than those in control cells (Fig. S3B’). Intriguingly, in *yki*^*3S/A*^ cells forming protrusions, Mys signals disappeared from the basal and the lateral sides of the cell body and accumulated at the protrusions (Fig. S3B’’, arrows, cell body; arrowheads, protrusion). We also assessed the subcellular distribution of Multiple edematous wings (Mew)–an α-subunit of *Drosophila* integrin. Mew signals were not readily detectable in control *esg*^+^ cells, probably due to low expression levels (Fig. S3C). In contrast, Mew signals were clearly visible at the basal and lateral sides in the majority of *yki*^*3S/A*^ cells, indicating that expression of *yki*^*3S/A*^ increased Mew (Fig. S3C’). Notably, in *yki*^*3S/A*^ cells forming protrusions, Mew signals were detected mainly at the protrusions (Fig. S3C’’, arrows, cell body; arrowheads, protrusion). Additionally, we checked the subcellular localization of Talin—a cytoplasmic protein that links integrins to the actin cytoskeleton. In control *esg*^*+*^ and most of the *yki*^*3S/A*^ cells, Talin signals were predominantly detected at the lateral and the basal sides (Fig. 1H and H’). In contrast, in *yki*^*3S/A*^ cells forming protrusions, strong Talin signals were observed at the protrusions but were negligible in the cell bodies (Fig. 1H’’, arrows, cell body; arrowheads, protrusion).

Taken together, these results show that only a portion of *yki*^*3S/A*^ cells can produce large protrusions across the VM that are enriched for focal adhesion components, again suggesting that only a subpopulation of *yki*^*3S/A*^ cells are migratory. The strong presence of Arm at cell-cell junctions of *yki*^*3S/A*^ cells indicates that most *yki*^*3S/A*^ cells are not migratory (Fig. 1A). Considering the assembly of focal adhesion components at the protrusions of *yki*^*3S/A*^ cells, these protrusions might serve as attachments for pulling out the cell bodies for cell dissemination.

### *yki*^*3S/A*^ tumors accumulate aberrantly heterogeneous cells in the EC lineage

Given that the phenotypes associated with invasive cell behavior were only observed in a portion of *yki*^*3S/A*^ cells, we speculated that *yki*^*3S/A*^ cells were not a homogeneous population. Additionally, *yki*^*3S/A*^ cells showing the invasive phenotypes were generally larger in size compared to control *esg*^*+*^ cells; their nuclei were also larger. During differentiation of EBs into ECs, EBs increase their size and ploidy. In contrast, EEs are generally smaller. Therefore, we hypothesized that measuring their nuclear size might be a way to look into the heterogeneity of *yki*^*3S/A*^ cells. In control midguts, *esg*^*ts*^-driven GFP marks ISCs and EBs, which are *esg*^*+*^ cells; ECs and EEs are not *esg*^*+*^ or *GFP*^*+*^. Likewise, *esg*^*ts*^-driven GFP marks *yki*^*3S/A*^ cells. The nuclear size measurement of *esg*^*+*^ cells in the control posterior midguts revealed three peaks (Fig. 2A, indicated with brackets and designated as I, II, and III). The main peak (I) was detected at around 11-15 μm, followed by a nuclei population (II) at around 19-23 μm (Fig. 2A). Additionally, relatively small nuclei formed a small peak (III) at around 6-9 μm (Fig. 2A). To understand how ISCs and EBs contribute to the overall distribution of *esg*^*+*^ nuclei, we also quantified the nuclear size of cells marked by *Dl*^*+*^—an ISC marker— or *Su(H)GBE*^*+*^—an EB marker. Interestingly, the *Su(H)GBE*^*+*^ nuclei size distribution showed a major peak at 11-15 μm and a lagging population at approximately 19-23 μm (Fig. 2A’, shown by brackets), which were reminiscent of the peaks I and II in the *esg*^*+*^ nuclear size distribution. The nuclear size distribution of *Dl*^*+*^ cells showed two peaks at 6-9 μm and 11-15 μm (Fig. 2A’’, indicated by brackets), which could be overlayed with the peaks III and I in the *esg*^*+*^ nuclear size distribution, respectively. These data suggest that both ISCs and EBs can contribute to the peak I while EBs and ISCs mainly contribute to the *esg*^*+*^ peak I and the *esg*^*+*^ peak III, respectively. We also measured the nuclear size of *esg*^*-*^ cells, which revealed two well-separated populations (Fig. S4A, arrows). Measuring the nuclear size of EEs, which are marked by an EE marker *prospero* (*pros*), showed a population of small nuclei (Fig. S4B) reminiscent of the *esg*^*-*^ small nuclei population at 6-9 μm. Thus, we concluded that the large *esg*^*-*^ nuclei population represented ECs (Fig. S4A). Interestingly, the *esg*^*+*^ peak III overlapped with the distribution of EE nuclei, which raised the possibility that the *esg*^*+*^ peak III might represent a population of *esg*^*+*^ cells in the EE lineage. Notably, the lagging *Su(H)GBE*^*+*^ population can be placed between the *esg*^*+*^ peak I and the EC nuclei peak at around 30 μm, suggesting that the lagging *esg*^*+*^ and *Su(H)GBE*^*+*^ populations might represent differentiating EBs. These analyses demonstrate that measuring nuclear size can be a way to assess the heterogeneity of midgut epithelial cells.

**Figure 2.**
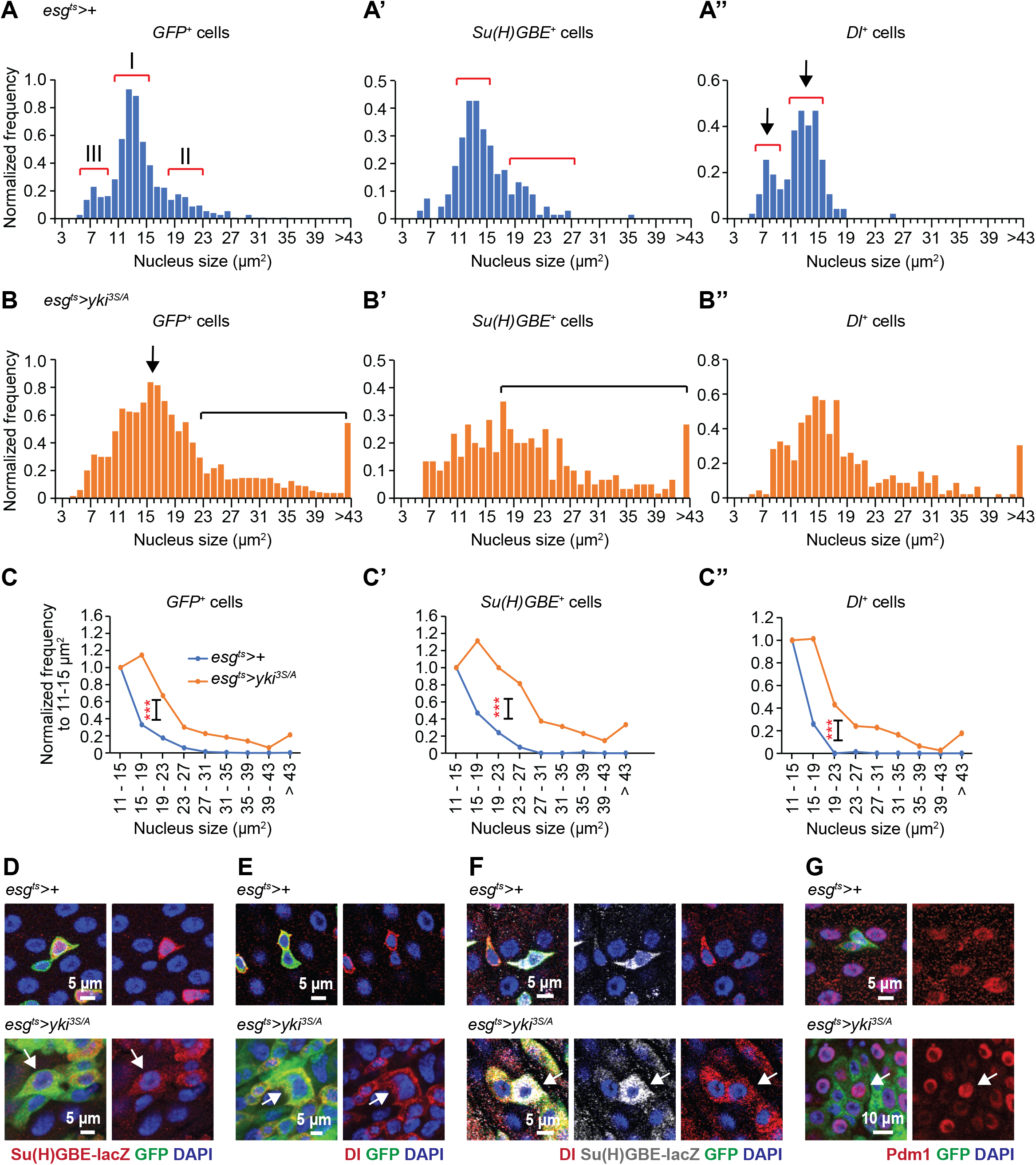
The EC differentiation program is aberrantly recapitulated in *yki*^*3S/A*^ midgut tumors, leading to generation of heterogeneous types of *yki*^*3S/A*^ cells. **A-A’’**. Nuclear size distribution of *GFP*^*+*^ cells (**A**), *Su(H)GBE*^*+*^ cells (**A’**), and *Dl*^*+*^ cells (**A’’**) in control (*esg*^*ts*^*>+*) midguts at day 6 of transgene induction. Numbered red brackets in (**A**) indicate three different cell populations described in the text. **B**. Nuclear size distribution of GFP^+^ cells (**B**), *Su(H)GBE*^*+*^ cells (**B’**), and *Dl*^*+*^ cells (**B’’**) in *esg*^*ts*^ >*yki*^*3S/A*^ midguts at day 6 of transgene induction. **C-C’’** Normalized Nuclear size distribution. Transgenes were induced for 6 days. The data is binned by 4 μm starting from 11 μm. Normalized frequency shows the population sizes relative to the population at 11-15 μm. *GFP*^*+*^ cells (**C**), N=691 (*esg*^*ts*^*>+*) and N=1869 (*esg*^*ts*^*>yki*^*3S/A*^); *Su(H)GBE*^*+*^ cells (**C’’**), N=179 (*esg*^*ts*^*>+*) and N=265 (*esg*^*ts*^*>yki*^*3S/A*^); *Dl*^*+*^ cells (**C’’**), N=103 (*esg*^*ts*^*>+*) and N=264 (*esg*^*ts*^*>yki*^*3S/A*^). ***P<0.001, chi-square test. **D**. *Su(H)GBE-lacZ* staining of posterior midguts at day 6. GFP marks *esg*^*+*^ cells (green). Arrow indicates a representative large *Su(H)GBE*^*+*^, *yki*^*3S/A*^ cell. **E**. Dl staining of posterior midguts at day 6. Arrow points to a representative extremely large *Dl*^*+*^, *yki*^*3S/A*^ cell. **F**. Dl and *Su(H)GBE-lacZ* co-staining of midguts at day 6. Arrow indicates a representative *Dl*^*+*^, *Su(H)GBE*^*+*^, *yki*^*3S/A*^ cell. **G**. Pdm1 staining of posterior midgut at day 6. Arrow points a representative *GFP*^*+*^, *Pdm1*^*+*^, *yki*^*3S/A*^ cells.

Given these results, we attempted to describe the population of *yki*^*3S/A*^ cells by measuring their nuclear size. As expression of *yki*^*3S/A*^ with *esg*^*ts*^ significantly increased cell division, significantly more *GFP*^*+*^ cells (11.83±0.35 cells per 50 μm x 50 μm region) were detected from a selected area in *yki*^*3S/A*^ midguts compared to control midguts (5.65±0.17 cells per 50 μm x 50 μm region). Although *yki*^*3S/A*^ nuclei were also populated at 11-15 μm, we observed a peak afterward (Fig. 2B, arrow) and a large lagging population of nuclei (Fig. 2B, bracket). Assuming that the population at 11-15 μm represents the majority of homeostatic ISCs and EBs, we decided to use the 11-15 μm population as an internal reference to bin the data by 4 μm, starting from 11 μm, to calculate the population sizes relative to the population at 11-15 μm in each genotype (Fig. 2C-C’’). This revealed a clear increase in the lagging *esg*^*+*^ nuclei population in *yki*^*3S/A*^ midguts compared to control midguts (Fig. 2C). Additionally, we observed an increase in the lagging *Su(H)GBE*^*+*^, *yki*^*3S/A*^ nuclei population (>15 μm), which might represent differentiating EB-like cells (Fig. 2B’ and C’). Intriguingly, large *Dl*^*+*^ nuclei (>15 μm) were more abundant in *yki*^*3S/A*^ midguts (Fig. 2B’’ and C’’). Therefore, accumulation of both large *Su(H)GBE*^*+*^ and *Dl*^*+*^ nuclei accounts for the increase in the lagging population of *yki*^*3S/A*^ nuclei. Unexpectedly, a significant portion of *Su(H)GBE*^*+*^, *yki*^*3S/A*^ nuclei were determined to be even larger than normal EC nuclei (Fig. 2B’ and D). Similarly, we also detected *Dl*^*+*^, *yki*^*3S/A*^ nuclei even larger than normal EC nuclei (Fig. 2B’’ and E). Strikingly, we found that a subpopulation of the *Su(H)GBE*^*+*^, *yki*^*3S/A*^ cells with large nuclei were also *Dl*^*+*^ (Fig. 2F), suggesting that these *Dl*^*+*^, *Su(H)GBE*^*+*^ cells might recapitulate the characteristics of ISCs. It appeared that a portion of *yki*^*3S/A*^ cells could undergo terminal differentiation as Pdm1 signals were detected in a portion of large *GFP*^*+*^ cells (Fig. 2G). Previous studies have shown that driving ISC differentiation towards ECs causes a reduction in EE cells (Beebe et al., 2010; Zeng and Hou, 2015). Indeed, EE cells were less frequently detected in regions where *yki*^*3S/A*^ cells were populated, implying that EE differentiation is reduced by the expression of *yki*^*3S/A*^ (Fig. S5). These observations suggest that *yki*^*3S/A*^ midguts still maintain the EC differentiation program even though it is not normal, resulting in an accumulation of a variety of cell populations, including aberrantly differentiating EB-like cells.

### Blocking the EC differentiation program in *yki*^*3S/A*^ tumors halts cell dissemination and abrogates JNK activation and Mmp1 expression in *yki*^*3S/A*^ cells

Given the accumulation of differentiating EB-like cells in *yki*^*3S/A*^ tumors, we decided to test the role of the EC differentiation program in dissemination of *yki*^*3S/A*^ cells. Notch signaling triggers the generation of EBs and the subsequent differentiation of EBs into ECs (Micchelli and Perrimon, 2006). If *yki*^*3S/A*^ cells use the same mechanism to generate the EB-like cells, inhibition of Notch signaling should significantly impact their differentiation process, which may result in an accumulation of ISC-like *yki*^*3S/A*^ cells. Thus, we expressed dominant-negative Notch (N^DN^) in *yki*^*3S/A*^ cells and then assessed the nuclear size distribution to gain insights into the changes in *yki*^*3S/A*^ cell populations. We found that *yki*^*3S/A*^, *N*^*DN*^ tumors grew as big as *yki*^*3S/A*^ tumors at day 8 of transgene induction (Fig. 3A). However, expression of N^DN^ in *yki*^*3S/A*^ cells resulted in a reduction in the lagging large nucleus population (Fig. 3B’ and C). In particular, most of the extremely large nuclei disappeared when N^DN^ and *yki*^*3S/A*^ were expressed together with *esg*^*ts*^ (Fig. 3B’ and C). Thus, *yki*^*3S/A*^, *N*^*DN*^ cells appeared to be more homogeneous than *yki*^*3S/A*^ cells and were positive for Dl (Fig. 3B-D), indicating that inhibition of Notch signaling in *yki*^*3S/A*^ cells efficiently blocked their differentiation and enriched ISC-like *yki*^*3S/A*^ cells. Of significance, expression of N^DN^ almost completely abolished dissemination of *yki*^*3S/A*^ cells at day 8 of transgene expression (Fig. 3E). Since a subpopulation of *yki*^*3S/A*^ cells showed the phenotypes associated with invasive cell behavior, we tested how blocking the generation of cells in the EC lineage affected Mmp1 expression and JNK signaling in *yki*^*3S/A*^, *N*^*DN*^ tumors. As we described previously, *puc-lacZ* and Mmp1 signals were detected in a portion of larger *yki*^*3S/A*^ cells and most of ECs in *yki*^*3S/A*^ midguts (Fig. 3F and G). In contrast, in *yki*^*3S/A*^, *N*^*DN*^ midguts, *puc-lacZ* and Mmp1 signals disappeared from *yki*^*3S/A*^, *N*^*DN*^ cells and were detected exclusively in ECs (*GFP*^*-*^ cells) (Fig. 3F and G). Taken together, these results demonstrate that blocking the EC differentiation program in *yki*^*3S/A*^ tumors is sufficient to attenuate the invasive behavior of *yki*^*3S/A*^ cells.

**Figure 3.**
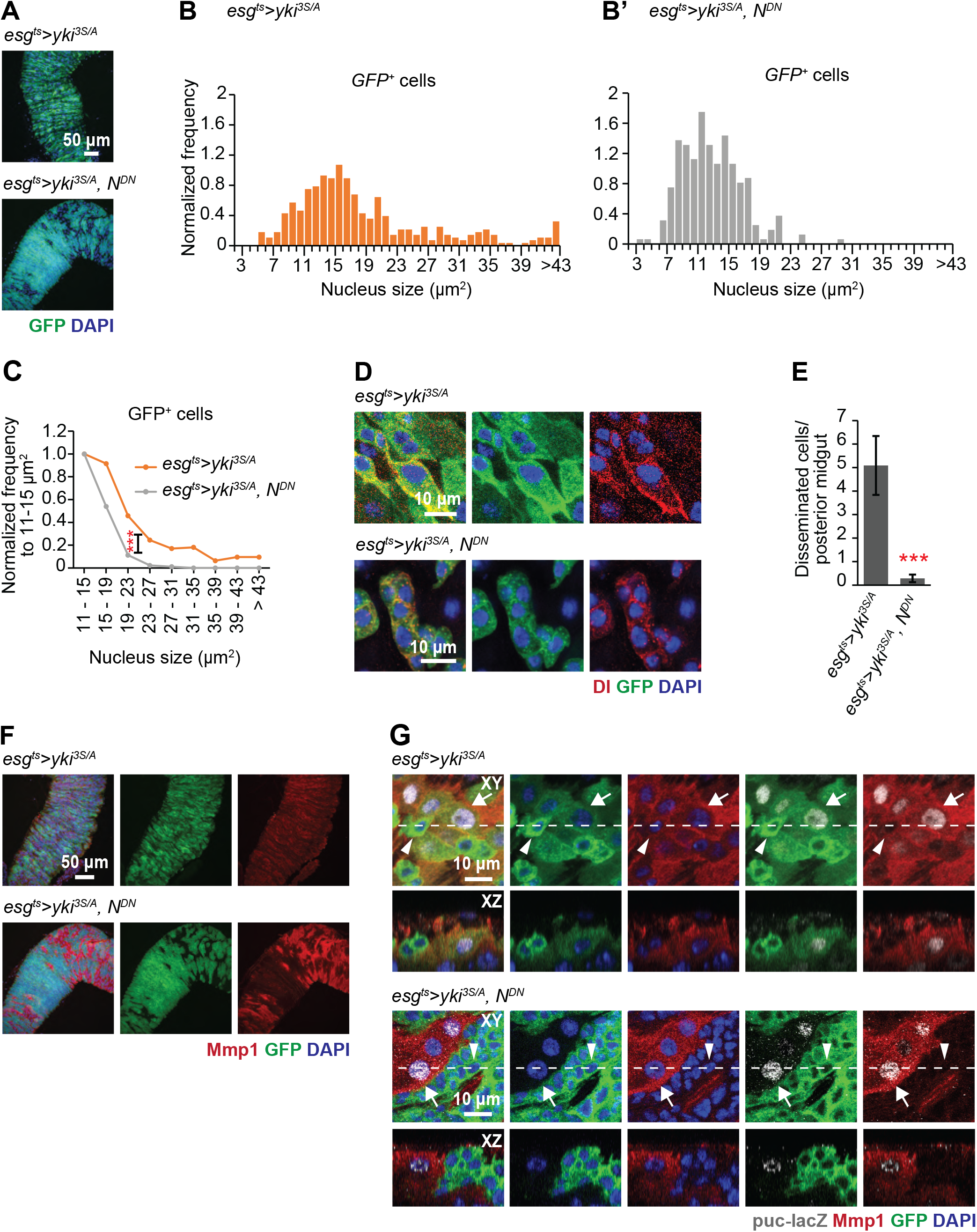
Arresting *yki*^*3S/A*^ cells in the ISC-like state suppresses Mmp1 expression in *yki*^*3S/A*^ cells and cell dissemination. **A**. Representative images of *esg*^*ts*^*>yki*^*3S/A*^ and *esg*^*ts*^ >*yki*^*3S/A*^, *N*^*DN*^ midgut tumors. Transgenes were induced for 8 days. Tumor cells are marked by GFP (green). **B-B’**. Nucleus size distribution of *GFP*^*+*^ cells in *esg*^*ts*^>*yki*^*3S/A*^ midgut (**B**) and *esg*^*ts*^>*yki*^*3S/A*^, *N*^*DN*^ midguts (B’) at day 6. **C**. Normalized N ㅜ uclear size distribution. Nucleus size distribution data shown in (B and B’) were binned by 4 μm starting from 11 μm, and then the binned values were normalized to the value from 11-15 μm^2^ in each genotype. N=303 (*esg*^*ts*^>*yki*^*3S/A*^) and N=150 (*esg*^*ts*^>*yki*^*3S/A*^, *N*^*DN*^) data points were analyzed. ***P<0.001, chi-square test. **D**. Representative images of *GFP*^*+*^ cells. D | signals are shown in red, and *yki*^*3S/A*^ and *yki*^*3S/A*^, *N*^*DN*^ cells are marked by GFP (green). Transgenes were induced for 6 days. **E**. Quantification of disseminated cells. Transgenes were induced for 8 days. N=22 (*esg*^*ts*^*>yki*^*3S/A*^) and N=14 (*esg*^*ts*^>*yki*^*3S/A*^, *N*^*DN*^) midguts. Mean±SEMs are shown. ***P<0.001, two-tailed unpaired Student’s t-test. **F**. Mmp1 staining (red) of posterior midguts after 8 days of transgene induction. **G**. Mmp1 and *puc-lacZ* co-staining of posterior midgut cells. Mmp1 signals are shown in red and *puc-lacZ* signals are shown in gray. Transgenes were induced for 6 days. Arrows show cells with Mmp1 signal and nuclear *puc-lacZ* signal, and arrowheads indicate cells lacking both Mmp1 and *puc-lacZ* signals.

### Blocking the EC differentiation program alters the tumor’s capacity to induce phenotypes associated with cachexia-like wasting

Tumors elicit various adverse effects on the host tissues and physiology in part by expressing secreted proteins (Argiles et al., 2014; Baracos et al., 2018; Figueroa-Clarevega and Bilder, 2015; Kim et al., 2021; Kwon et al., 2015; Petruzzelli and Wagner, 2016; Porporato, 2016; Song et al., 2019; Yeom et al., 2021). In particular, *yki*^*3S/A*^ midgut tumors express multiple secreted factors, which play key roles in inducing the phenotypes associated with cachexia-like wasting, such as organ degeneration, metabolic abnormalities, and reduced lifespan (Kim et al., 2021; Kwon et al., 2015; Song et al., 2019). Given the observation that blocking the EC differentiation program in *yki*^*3S/A*^ tumors suppressed the phenotypes associated with invasiveness, we assessed how expression of *N*^*DN*^ in *yki*^*3S/A*^ tumors altered the tumor’s propensity to induce the non-tumor-autonomous or systemic phenotypes. Blocking the EC differentiation program in *yki*^*3S/A*^ tumors significantly suppressed the ‘bloating syndrome’ phenotype— the manifestation of cachexia-like wasting—which is characterized by a bloated abdomen due to an increased circulating fluid (Fig. 4A, upper panels and B). Expression of N^DN^ also fully rescued ovary atrophy observed in flies harboring *yki*^*3S/A*^ midgut tumors (Fig. 4A, bottom panels and C). The adult visceral cavity is filled with amorphous fat body–the adipose tissue in *Drosophila*–, which makes the ventral side of the abdomen opaquely yellow-whiteish (Kwon et al., 2015). Since the fat body significantly degenerated in flies bearing *yki*^*3S/A*^ midgut tumors, the abdomen became translucent (Fig. 4A, middle panels). In contrast, the abdomen of flies bearing *yki*^*3S/A*^, *N*^*DN*^ midgut tumors mostly remained opaque (Fig. 4A, middle panels), indicating that expression of *N*^*DN*^ in *yki*^*3S/A*^ tumors suppressed fat body degeneration. These wasting phenotypes are associated with the expression of the secreted antagonist of *Drosophila* insulin-like peptides (Dilps), ImpL2 (Honegger et al., 2008; Kwon et al., 2015; Lee et al., 2021). ImpL2 expressed in *yki*^*3S/A*^ tumors induced a systemic reduction in insulin/IGF signaling, resulting in hyperglycemia (Kwon et al., 2015). Of note, expression of *N*^*DN*^ in *yki*^*3S/A*^ midgut tumors significantly rescued the hyperglycemia phenotype, which was assessed by measuring trehalose levels (Fig. 4D). However, expression of *N*^*DN*^ in *yki*^*3S/A*^ midgut tumors did not alleviate all the phenotypes associated with cachexia-like wasting. We assessed muscle degeneration in tumor-bearing flies by measuring their climbing defects. Interestingly, expression of *N*^*DN*^ in the tumors failed to rescue climbing defects associated with *yki*^*3S/A*^ tumors (Fig. 4E). Moreover, blocking the EC differentiation program in *yki*^*3S/A*^ midgut tumors further shortened the lifespan of the tumor-bearing flies (Fig. 4F and S6). These observations differentiate the roles of the ISC-like and the differentiating EB-like cells in inducing the phenotypes associated with cachexia-like wasting. Moreover, we show that the differentiation program recapitulated in *yki*^*3S/A*^ tumors plays a key role in shaping the tumor’s capacity to induce various non-tumor-autonomous phenotypes.

**Figure 4.**
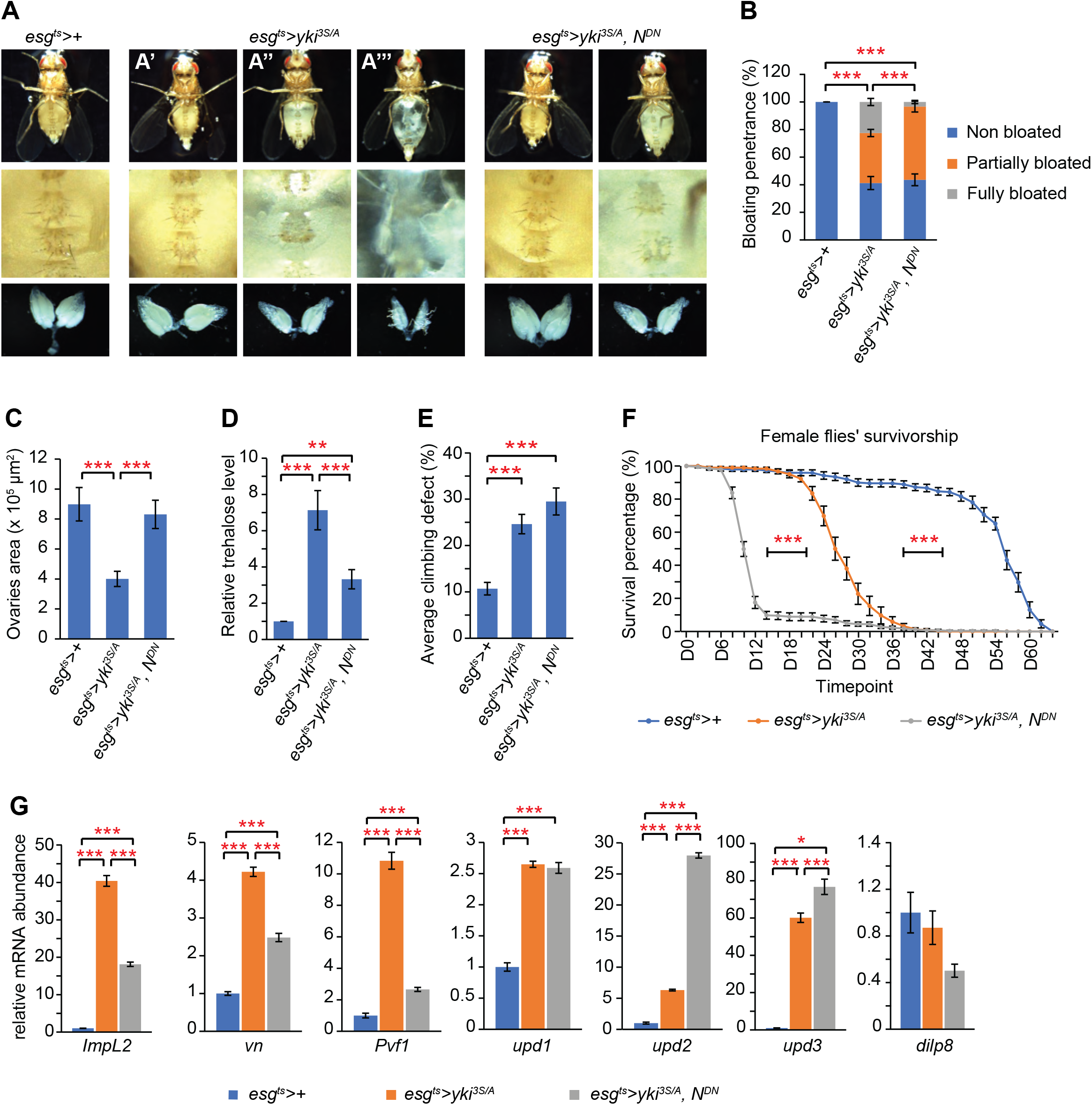
Blocking the EC differentiation program in *yki*^*3S/A*^ cells alters the phenotypes associated with cachexia-like wasting. **A**. Representative ventral views and ovary images. Upper panels, ventral views; middle panels, magnified views of the corresponding fly’s abdominal area; lower panels, images of the corresponding fly’s ovaries. The upper panels in *esg*^*ts*^*>yki*^*3S/A*^ show representative non-bloated (normal state) (**A’**), partially bloated (**A’’**), and fully bloated (**A’’**) fly’s abdominal views. Transgenes were induced for 8 days. **B**. Quantification of bloating syndrome penetrance. Transgenes were induced for 8 days. N=481 (*esg*^*ts*^*>+*), N=568 (*esg*^*ts*^*>yki*^*3S/A*^), and N=375 (*esg*^*ts*^>*yki*^*3S/A*^, *N*^*DN*^) animals. Mean±SEMs are shown. ***P<0.001, chi-square test. **C**. Quantification of ovaries size. Transgenes were induced for 8 days. N=19 (*esg*^*ts*^*>+*), N=20 (*esg*^*ts*^*>yki*^*3S/A*^), and N=17 (*esg*^*ts*^>*yki*^*3S/A*^, *N*^*DN*^) pairs of ovaries. Mean±SEMs are shown. ***P<0.001, two-tailed unpaired Student’s t-test. **D**. Relative trehalose levels in *esg*^*ts*^*>+, esg*^*ts*^*>yki*^*3S/A*^, *and esg*^*ts*^*>yki*^*3S/A*^, *N*^*DN*^ flies at day 8 of transgene induction. Mean±SEMs are shown. N=6 flies, 6 biological replicates. ***P<0.001, two-tailed unpaired Student’s t-test. **E**. Quantification of climbing defect in *esg*^*ts*^*>+, esg*^*ts*^*>yki*^*3S/A*^, and *esg*^*ts*^*>yki*^*3S/A*^, *N*^*DN*^ flies at day 8 of transgene induction. N=608 (*esg*^*ts*^*>+*), N=608 (*esg*^*ts*^*>yki*^*3S/A*^), and N=204 (*esg*^*ts*^>*yki*^*3S/A*^, *N*^*DN*^). Mean±SEMs are shown. ***P<0.001, two-tailed unpaired Student’s t-test. **F**. Survivorship in female flies. The experiment was performed at 29°C to maintain the expression of transgenes. *esg*^*ts*^*>+*, blue, N=159 flies in 17 replicates; *esg*^*ts*^*>yki*^*3S/A*^, orange, N=221 flies in 17 replicates; *esg*^*ts*^*>yki*^*3S/A*^, *N*^*DN*^, gray, N= 224 flies in 18 replicates. Mean±SEMs are shown. ***p < 0.001, log-rank test. **G**. Relative mRNA levels of various wasting factors at day 8 of transgene induction. N=10 female midguts for each genotype, 3 biological replicates. Mean±SEMs are shown. *P<0.05, **P<0.01, ***p < 0.001, two-tailed unpaired Student’s t-test.

Previous studies have identified several tumor-derived factors eliciting various non-tumor-autonomous phenotypes (Bilder et al., 2021; Figueroa-Clarevega and Bilder, 2015; Kim et al., 2021; Kwon et al., 2015; Liu et al., 2022; Lodge et al., 2021; Song et al., 2019; Yeom et al., 2021).Thus, we assessed the expression of various tumor-driven factors. Indeed, we found that blocking the EC differentiation program in *yki*^*3S/A*^ midgut tumors significantly impacted the expression of several tumor-derived factors. Previous studies have shown that tumor-derived ImpL2 and epithermal growth factors (EGFs), such as PDGF- and VEGF-related factor 1 (Pvf1) are required for inducing the wasting phenotypes, including ovary atrophy, muscle degeneration, hyperglycemia, and bloating syndrome (Figueroa-Clarevega and Bilder, 2015; Kwon et al., 2015; Song et al., 2019; Yeom et al., 2021). Notably, mRNA levels of *ImpL2* and *Pvf1* were significantly reduced in *yki*^*3S/A*^, *N*^*DN*^ tumors compared to *yki*^*3S/A*^ tumors. An EGF ligand, *vein* (*vn*) was shown to function locally to support *yki*^*3S/A*^ tumor growth (Song et al., 2019). *vn* mRNA levels were also reduced while tumor growth was not affected (Fig. 4G). Recently, Kim et al. showed that the *Drosophila* Interleukin-6 orthologs Unpaired 2 (Upd2) and Unpaired 3 (Upd3) derived from imaginal disc tumors caused an aberrant permeability of the blood-brain barrier (BBB), which was responsible for the significant reduction in the lifespan of the tumor-bearing flies (Kim et al., 2021). As the BBB was also shown to be compromised in the flies bearing *yki*^*3S/A*^ midgut tumors (Kim et al., 2021), we tested the expression of all three *unpaired* genes. *upd1* mRNA levels were similar in both tumors while *upd2 and upd3* mRNA levels were significantly increased in *yki*^*3S/A*^, *N*^*DN*^ tumors compared to *yki*^*3S/A*^ tumors (Fig. 4G). Yoem et al. has reported that eye tumors induced by an active *yki* allele (*yki*^*S168A*^) express a *Drosophila* homolog of the mammalian insulin-like 3 peptide (INSL3) Dilp8, which induces anorexia in flies (Yeom et al., 2021). However, *dilp8* mRNA levels were unaltered in *yki*^*3S/A*^ midgut tumors compared to control midguts (Fig. 4G), and expression of N^DN^ also did not significantly change *Dilp8* mRNA levels (Fig. 4G). These results indicate that blocking the EC differentiation program in *yki*^*3S/A*^ midgut tumors alters the expression of the tumor-derived factors, which are responsible for the adverse non-tumor-autonomous or systemic phenotypes.

Given the role of ImpL2 and Pvf1 in inducing various wasting phenotypes, the reduction in *ImpL2* and *Pvf1* expression might account for the suppression of bloating syndrome, ovary atrophy, fat body degeneration, and hyperglycemia in flies bearing *yki*^*3S/A*^, *N*^*DN*^ tumors. Additionally, the elevated expression of *upd2* could be a factor responsible for the further shortening of the lifespan in flies bearing *yki*^*3S/A*^, *N*^*DN*^ tumors as tumor-derived Upd2 impairs the BBB. Importantly, our observations indicate that ImpL2 is mainly expressed in *yki*^*3S/A*^ cells in the EC lineage even though ISC-like *yki*^*3S/A*^ cells also express ImpL2 (Fig. 4G). Depletion of *ImpL2* in *yki*^*3S/A*^, *N*^*DN*^ tumors further suppressed bloating syndrome and hyperglycemia (Fig. S7), indicating that residual ImpL2 expression in *yki*^*3S/A*^, *N*^*DN*^ tumors was responsible for the incomplete suppression of these phenotypes by the elimination of the *yki*^*3S/A*^ cells in the EC lineage. The significant increase in *upd2* mRNA expression in *yki*^*3S/A*^, *N*^*DN*^ tumors suggests that Upd2 is predominantly expressed in ISC-like *yki*^*3S/A*^ cells. Notably, depletion of *upd2* resulted in complete suppression of *yki*^*3S/A*^, *N*^*DN*^ tumor growth (Fig. S8), supporting that Upd2 is expressed in ISC-like *yki*^*3S/A*^ cells. Previous reports have shown that Upd2 is expressed in EBs and ECs during tissue maintenance and regeneration (Jiang et al., 2009; Zhai et al., 2015). Our results suggest a role for the ISC-like *yki*^*3S/A*^ cell-derived Upd2 in inducing tumor growth as well as non-tumor-autonomous phenotypes. Altogether, our findings suggest that tumor cell type-dependent expression of the tumor-derived factors could be the mechanism by which halting the EC differentiation program alters the tumor’s capacity to induce non-tumor-autonomous phenotypes.

## Discussion

We demonstrate that the EC differentiation program is aberrantly recapitulated in *yki*^*3S/A*^ midgut tumors, resulting in the generation of heterogeneous types of *yki*^*3S/A*^ cells. Based on the markers of ISCs and EBs, Dl and Su(H)GBE-lacZ, respectively, *yki*^*3S/A*^ cells could be divided into at least 3 different types: ISC-like (*Dl*^*+*^), EB-like (*Su(H)GBE*^*+*^), and ISC/EB-like (*Dl*^*+*^, *Su(H)GBE*^*+*^) (Fig. 2B-F). Although *Dl*^*+*^, *Su(H)GBE*^*+*^ cells were rarely detected in control midguts, a portion of *yki*^*3S/A*^ cells were *Dl*^*+*^, *Su(H)GBE*^*+*^ (Fig. 2F). We observed an accumulation of *yki*^*3S/A*^ cells with large nuclei, which were reminiscent of EBs undergoing differentiation into ECs (Fig. 2B’ and D). The EC cell marker Pdm1 was occasionally detected in large *Su(H)GBE*^*+*^ cells with reduced GFP signals. Thus, at least some *yki*^*3S/A*^ cells appeared to undergo terminal differentiation to generate ECs (Fig. 2G). Interestingly, some nuclei of *yki*^*3S/A*^ cells were even larger than the nuclei of normal EC cells, suggesting that a portion of *yki*^*3S/A*^ cells deviate from the normal EC lineage (Fig. 2B). In the midgut epithelium, ISCs by default generate ISCs, and activation of Notch signaling triggers the EC lineage by generating EBs and initiating the differentiation of EBs into ECs (Boumard and Bardin, 2021; Hou and Singh, 2017; Jiang et al., 2016; Li and Jasper, 2016; Micchelli and Perrimon, 2006; Miguel-Aliaga et al., 2018; Ohlstein and Spradling, 2006). Of importance, inhibiting Notch signaling in *yki*^*3S/A*^ tumors dramatically diminished the cellular heterogeneity by eliminating *yki*^*3S/A*^ cells resembling the cells in the EC lineage, resulting in an accumulation of ISC-like *yki*^*3S/A*^ cells (*Dl*^*+*^). Altogether, our study reveals the heterogeneity of *yki*^*3S/A*^ midgut tumor cells and elucidates the key role of the native EC differentiation program in generation of the cellular heterogeneity.

Recent studies have revealed that the developmental and/or differentiation programs that form and maintain the tissues of origin are recapitulated in cancers, such as glioblastoma, Wilms tumors, and non-small-cell lung cancer (Couturier et al., 2020; Fukuzawa et al., 2017; Goveia et al., 2020; Kim et al., 2020; Wu et al., 2021; Yeo et al., 2020). However, it remains unclear how the developmental and differentiation programs affect tumor progression and the tumor’s capacity to induce various non-tumor-autonomous phenotypes, such as cachexia. Our findings provide us with an opportunity to use *Drosophila* genetics to discriminate the roles of ISC-like tumor cells and differentiating tumor cells in tumor progression as well as inducing the phenotypes associated with cachexia-like wasting. *yki*^*3S/A*^ cells co-opting the EC differentiation program significantly accumulate, and inhibition of Notch signaling eliminates these differentiating *yki*^*3S/A*^ cells, leading to a formation of tumors comprised of ISC-like *yki*^*3S/A*^ cells. Of note, *yki*^*3S/A*^, *N*^*DN*^ tumors can still robustly grow, indicating that the differentiating tumor cells are essentially dispensable for tumor formation and growth. Interestingly, eliminating the differentiating tumor cells almost completely suppresses tumor cell dissemination and attenuates the phenotypes associated with invasive cell behavior. Thus, our study reveals an unexpected role of the differentiating tumor cells in the invasion process. It has been shown that Esg is expressed more in the EBs committed to differentiation than ISCs (Antonello et al., 2015; Korzelius et al., 2014). The committed EBs acquire mesenchymal characteristics, such as polarized shape and invasive properties (Antonello et al., 2015). This raises an interesting possibility that the intrinsically invasive properties of the EBs undergoing differentiation could be the origin of the invasiveness of *yki*^*3S/A*^ cells. We speculate that the accumulation of *yki*^*3S/A*^ cells resembling the intrinsically-invasive EB pool might result in cell dissemination. Coordination of cell differentiation and migration is a widely spread phenomenon in the development and tissue maintenance (Aalto et al., 2021; Antonello et al., 2015; Arvidsson et al., 2002; Montell, 2003; Parent et al., 2002; Peercy and Starz-Gaiano, 2020; Vasilyev et al., 2009). Therefore, we propose that an accumulation of differentiating tumor cells induced by the developmental and differentiation programs recapitulated in tumors could be a mechanism by which tumors acquire invasiveness.

Our study elucidates the differential roles of ISC-like and differentiating tumor cells in inducing the phenotypes associated with cachexia-like wasting. Depletion of the differentiating tumor cells by expressing N^DN^ significantly rescues an array of the phenotypes associated with cachexia-like wasting: ovary atrophy, fat body degeneration, hyperglycemia, and bloating syndrome. However, blocking cell differentiation does not rescue muscle degeneration and further shortens the lifespan. These observations demonstrate the distinct roles ISC-like and differentiating tumor cells play in shaping the tumor’s capacity to induce various non-tumor-autonomous phenotypes. Changing the tumor-cell population by blocking cell differentiation causes significant alterations in the expression of the tumor-derived wasting factors. Depletion of the differentiating tumor cells greatly reduces *ImpL2* mRNA expression in the tumors, and depletion of *ImpL2* in *yki*^*3S/A*^, *N*^*DN*^ tumors further suppressed hyperglycemia and bloating syndrome, indicating that *ImpL2* is predominantly expressed in the differentiating tumor cells. In contrast, depletion of the differentiating tumor cells significantly increases *upd2* mRNA levels, and depletion of *upd2* in *yki*^*3S/A*^, *N*^*DN*^ cells significantly suppressed tumor growth, indicating that Upd2 is primarily expressed in ISC-like tumor cells. Given the role of Upd2 in impairing the normal BBB function, the increase in Upd2 expression in *yki*^*3S/A*^, *N*^*DN*^ tumors could account for the reduction in the lifespan of the tumor-bearing flies (Kim et al., 2021). Thus, these observations indicate that these tumor-derived factors are differentially expressed in ISC-like and differentiating tumor cells, which could be the basis for the alteration of non-tumor autonomous phenotypes upon reducing cellular heterogeneity. Together these results show that tumor cell heterogeneity is a key factor that shapes the tumor’s capacity to induce various non-tumor-autonomous and systemic phenotypes. Furthermore, our study provides mechanistic insight into how eliminating a tumor cell population can alter the tumor’s capacity to induce non-tumor-autonomous phenotypes.

Simple genetic manipulations, such as the expression of an oncogene and the depletion of a tumor suppressor, induce hyperplasia in various tissues in *Drosophila*. These hyperplasias have been used as so-called *Drosophila* “tumor” models, which has led to the discovery of various fundamental mechanisms underlying the growth of normal tissues as well as cancers. Notably, *Drosophila* tumors can also induce various phenotypes reminiscent of those observed in advanced cancer patients. Since hyperplasias are normally induced for a few days, it is unlikely that these tumors acquire additional genetic alterations, which can confer new properties to the tumors. Therefore, the ability of tumors to induce certain phenotypes cannot be entirely dependent on the gain of an oncogene or loss of a tumor suppressor. Our study provides insights into how the native cell differentiation program can contribute to the tumor’s capacity to induce various phenotypes. Moreover, our study raises the possibility of manipulating the advanced cancer phenotypes by altering the developmental and differentiation programs co-opted in tumors. Thus, it would be of interest to address how the developmental and differentiation programs recapitulated in cancers contribute to the induction of complications associated with cancers, which might eventually lead to a strategy to treat cancer cachexia.

## Supporting information

Supplemental figures (Revision)

## Acknowledgments

We thank Dr. Page-McCaw for sharing Mmp1 transgenic fly lines and Dr. Cai for allocating anti-Pdm1 antibody. This work was supported by R35GM128752 to Y.K. from the National Institutes of Health.

## Author contributions

I.P. and Y.K. designed experiments, analyzed data, and wrote the manuscript. I.P. performed experiments. J.L. made the observation that allowed us to initiate the project.

## Methods

### Fly genetics and husbandry

Fly crosses were maintained in vials with standard cornmeal-agar medium and kept at 18°C throughout development and adulthood until ready for temperature-dependent induction. For all experiments, except for climbing and lifespan assays, zero to three-day-old female flies were collected and were shifted to 29°C to induce transgene expression for the indicated number of days prior to dissection. During the 29°C incubation, flies were transferred to fresh food vials every 2 days. To manipulate ISCs and EBs, we used *esg-GAL4, tub-GAL80*^*ts*^, *UAS-GFP* (referred as *esg*^*ts*^). Fly strains used in this study included: *UAS-yki*^*3S/A*^ (*w\**;; *UAS-yki*.*S111A*.*S168A*.*S250A*.*V5*) (BDSC #28817), *Su(H)GBE-lacZ* (BDSC #83352), *UAS-Mmp1 RNAi* (BDSC #31489), *puc-lacZ* (laboratory stock), *UAS-Notch*^*DN*^ (laboratory stock), and *UAS-Mmp1*^*DN Pro-pex*^ (a gift from the Page-McCaw Lab).

### Antibodies and immunofluorescence imaging

Immunostainings performed in this study used the following primary antibodies: anti-GFP antibody, Alexa Fluor® 488 (1:1000; Thermo Fisher Scientific, A-21311; rabbit), anti-Armadillo antibody (1:1000; DSHB, N2 7A1; mouse), anti-laminin B1 antibody (1:1000; Abcam, ab47650; rabbit), anti-Mmp1 antibody (1:100; DSHB, 3B8D12; mouse), anti-Mys antibody (1:1000; DSHB, CF.6G11; mouse), anti-Mew antibody (1:300; DSHB, DK.1A4; mouse), anti-Talin antibody (1:1000; DSHB, A22A; mouse), anti-Delta antibody (1:1000; DSHB, C594.9B; mouse), anti-Pdm1 antibody (1:500; a gift from the Cai Lab; rabbit), anti-β Galactosidase antibody (1:1000; Cappel, 55976; rabbit and 1:1000; DSHB, 40-1a-s; mouse), anti-Prospero antibody (1:1000; DSHB, MR1A; mouse). All secondary antibodies were obtained from Thermo Fisher Scientific: anti-rabbit IgGs conjugated to Alexa Flour® 594 (1:1000; A-11012; goat), anti-mouse IgGs conjugated to Alexa Flour® 594 (1:1000; A-11005; goat), anti-rabbit IgGs conjugated to Alexa Flour® 647 (1:1000; A-21244; goat), and anti-mouse IgGs conjugated to Alexa Flour® 647 (1:1000; A-21244; goat). Filamentous actin was stained with phalloidin conjugated to Alexa Fluor® 594 or 647 (1:1000; Thermo Fisher Scientific, A-12381, A-22287, respectively). Nuclei were stained with DAPI (1:2000; Sigma, D9542).

To remove auto-fluorescing remnants from the midguts, we fed flies 4% sucrose for ∼4 hr prior to dissection. Midgut samples were dissected in PBS, fixed in 4% paraformaldehyde (PFA) (Electron Microscopy Sciences, RT15710) diluted in PBS for 20 min, and then washed three times with PBST (PBS supplemented with 0.2% Triton X-100) with 5 min intervals. For permeabilization and blocking, we incubated tissue samples in blocking buffer (PBST supplemented with 5% normal goat serum) for 1 hr at room temperature. The tissue samples were incubated with primary antibodies in the blocking buffer overnight at 4°C. The samples were washed three times with PBST and then incubated in secondary antibodies for 2 hr at room temperature. Stained midguts were washed three times with PBST and preserved in Vectashield (Vector Laboratories, H-1000). Fluorescence images were acquired using a Leica SP8 laser scanning confocal microscope with 40×/1.25 oil objective lens. NIH ImageJ software was used for further adjustment and assembly of the acquired images.

### Quantification of Armadillo signals

To measure the distribution and fluorescence intensity of Arm signals, we created a z-projection of the topmost layer of the front epithelium leaflet for each 290.91 μm × 290.91 μm z-stacks posterior midgut image. We drew a line from the outside of a cell across the cell at multiple individual cells. Mean gray values of the red channel (Arm signal) were collected using the Plot Profile function on NIH ImageJ software, giving a list of mean gray values at each distance point along the drawn line. These distance points were then adjusted to account for the line length differences among cells. Because Arm signals are predominantly at the membrane, the points where the line intersects the membrane would give the highest signal values. The distance point where the line first intersects the membrane was then calibrated as 0 μm, resulting in the distance points outside the cell as -X μm and distance points across inside of the cells as +X μm.

### Quantification of disseminated cells

Disseminated cells quantified in this study were defined as *GFP*^*+*^ and *DAPI*^*+*^ cells residing more basally than the visceral muscle (labeled with phalloidin) of the front epithelial leaflet. A series of z-stack images were taken to capture the posterior midguts using confocal microscopy. To determine cell positions respective to the visceral muscle layer, we reconstituted orthogonal views from the z-stacks images. The total number of disseminated cells was quantified in 290.91 μm × 290.91 μm confocal microscope fields.

### Quantification of Mmp1 intensity

To measure the fluorescence intensity of Mmp1, we imaged the posterior midgut epithelium using confocal microscopy and generated z-projections. Mean gray values of the red channel (Mmp1 signal) were collected using 2.84 μm × 2.84 μm fields for ten random *GFP*^*+*^ cells and ten random *GFP*^*-*^ cells (ECs) per intestine using NIH ImageJ software. The values were then subtracted by the background value, which was obtained by measuring the mean gray value of the outside area surrounding the tissue. All intensity values were normalized to the intensity value of *GFP*^*+*^ cells in *esg*^*ts*^ intestine.

### Nucleus size measurement analysis

To measure nucleus size, we created a z-projection of a 50 μm × 50 μm region capturing the topmost epithelial leaflet from a 290.91 μm × 290.91 μm posterior midgut image. We generated an RGB image of the DAPI channel (nuclei channel) only, made the image to an 8-bit type image, and converted them into a binary image. We applied a feature called Watershed to separate nuclear clusters due to cell crowding. The area of each nucleus was measured by subjecting the final binary image to Analyze Particles feature on NIH ImageJ software. All area measurements were filtered manually to exclude artifact quantifications by comparing the binary image to the original 50 μm × 50 μm RGB image. At least four-50 μm × 50 μm-regions were created and analyzed for each intestine to cover the span of the posterior midgut.

### Quantification of Prospero-positive cells

To determine the number of EE cells, we dissected midguts and stained them with DAPI and anti-Prospero antibody. *pros*^*+*^ nuclei were counted using a 100 μm × 100 μm field from posterior-end and anterior-end of *esg*^*ts*^ and *yki*^*3S/A*^ posterior midgut region.

### Measurement of trehalose level

To prepare fly lysates for trehalose assays, we homogenized six female flies of each genotype in 400 μL PBST, heated the lysate at 70°C for 5 min, centrifuged the samples at 14,000 rpm for 10 min, and collected the supernatant. Whole-body trehalose levels were measured using a trehalose assay kit (Neogen; K-TREH) according to the manufacturer’s protocol. The final value of trehalose levels was normalized to number of flies and then to trehalose level in control flies to obtain relative trehalose levels for each genotype.

### Quantification of flies climbing activity assay

To assess climbing activity, we used flies at 14 days post induction. We transferred them into vials with a 2-cm mark from the food surface with a maximum capacity of 10 flies per vial. To record the climbing ability, we tapped down a vial for five times to bring the flies to the bottom and recorded the number of flies that climbed to pass the 2-cm mark in 3 seconds. Flies with a climbing defect were defined as those who were not able to pass the mark in 3 seconds. Climbing activity assay was recorded 3 times for each vial.

### Drosophila lifespan assay

We collected adult female and male flies in separate vials with less than 25 individuals per vial and kept them at 29 °C incubator for transgenes induction. We transferred flies into fresh food vials and recorded deaths every two days.

### Quantitative RT-PCR

We isolated total RNA from 20 adult female midguts or six female thoraces at D8 of induction with TRIzol (Invitrogen, Cat# 15596026). We used 1 μg of RNA to produce cDNA with iScript™ Reverse Transcription Supermix (Bio-Rad, Cat#1725120). The cDNA was subjected to quantitative real-time PCR with iTaq™ Universal SYBR Green Supermix (Bio-Rad, Cat#1708840) and CFX-96 (Bio-Rad). The fold-changes in RNA transcript levels were normalized against RpL32 gene, and further normalized to the control genotype for relative mRNA expression levels. Primer sequences are the following: Pvf1, CTGTCCGTGTCCGCTGAG, CTCGCCGGACACATCGTAG; vn, GAACGCAGAGGTCACGAAGA, GAGCGCACTATTAGCTCGGA; dilp8, GGACGGACGGGTTAACCATT, CATCAGGCAACAGACTCCGA; ImpL2, AAGAGCCGTGGACCTGGTA, TTGGTGAACTTGAGCCAGTCG; upd1, CCTACTCGTCCTGCTCCTTG, TGCGATAGTCGATCCAGTTG; upd2, CATCGTCATCCTCATCATCG, ATGTTCCGCAAGTTTTCGAG; upd3, AAATTCGACAAAGTCGCCTG, TTCCACTGGATTCCTGGTTC; RP49, GCTAAGCTGTCGCACAAATG, GTTCGATCCGTAACCGATGT.

### Statistics and reproducibility

We performed all statistical analyses using Microsoft Excel. Levels of significance are depicted by asterisks in the figures: *P<0.05, **P<0.01, ***P<0.001. Sample sizes were chosen empirically based on the observed effects and listed in the figure legends.

## References

Aalto, A., Olguin-Olguin, A., and Raz, E. (2021). Zebrafish Primordial Germ Cell Migration. Front Cell Dev Biol 9, 684460.

Antonello, Z.A., Reiff, T., Ballesta-Illan, E., and Dominguez, M. (2015). Robust intestinal homeostasis relies on cellular plasticity in enteroblasts mediated by miR-8–Escargot switch. The EMBO Journal 34, 2025–2041.

Apidianakis, Y., Pitsouli, C., Perrimon, N., and Rahme, L. (2009). Synergy between bacterial infection and genetic predisposition in intestinal dysplasia. Proc Natl Acad Sci U S A 106, 20883–20888.

Argiles, J.M., Busquets, S., Stemmler, B., and Lopez-Soriano, F.J. (2014). Cancer cachexia: understanding the molecular basis. Nat Rev Cancer 14, 754–762.

Arvidsson, A., Collin, T., Kirik, D., Kokaia, Z., and Lindvall, O. (2002). Neuronal replacement from endogenous precursors in the adult brain after stroke. Nat Med 8, 963–970.

Baracos, V.E., Martin, L., Korc, M., Guttridge, D.C., and Fearon, K.C.H. (2018). Cancer-associated cachexia. Nat Rev Dis Primers 4, 17105.

Beebe, K., Lee, W.C., and Micchelli, C.A. (2010). JAK/STAT signaling coordinates stem cell proliferation and multilineage differentiation in the Drosophila intestinal stem cell lineage. Developmental Biology 338, 28–37.

Bilder, D., Ong, K., Hsi, T.C., Adiga, K., and Kim, J. (2021). Tumour-host interactions through the lens of Drosophila. Nat Rev Cancer 21, 687–700.

Biteau, B., and Jasper, H. (2014). Slit/Robo signaling regulates cell fate decisions in the intestinal stem cell lineage of Drosophila. Cell Rep 7, 1867–1875.

Borczuk, A.C., Gorenstein, L., Walter, K.L., Assaad, A.A., Wang, L., and Powell, C.A. (2003). Non-small-cell lung cancer molecular signatures recapitulate lung developmental pathways. Am J Pathol 163, 1949–1960.

Boumard, B., and Bardin, A.J. (2021). An amuse-bouche of stem cell regulation: Underlying principles and mechanisms from adult Drosophila intestinal stem cells. Curr Opin Cell Biol 73, 58–68.

Chen, K., Huang, Y.H., and Chen, J.L. (2013). Understanding and targeting cancer stem cells: Therapeutic implications and challenges. In Acta Pharmacologica Sinica, pp. 732–740.

Cook, J.H., Melloni, G.E.M., Gulhan, D.C., Park, P.J., and Haigis, K.M. (2021). The origins and genetic interactions of KRAS mutations are allele-and tissue-specific. Nature Communications 12.

Couturier, C.P., Ayyadhury, S., Le, P.U., Nadaf, J., Monlong, J., Riva, G., Allache, R., Baig, S., Yan, X., Bourgey, M., et al. (2020). Single-cell RNA-seq reveals that glioblastoma recapitulates a normal neurodevelopmental hierarchy. Nature Communications 11.

da Silva-Diz, V., Lorenzo-Sanz, L., Bernat-Peguera, A., Lopez-Cerda, M., and Muñoz, P. (2018). Cancer cell plasticity: Impact on tumor progression and therapy response. In Seminars in Cancer Biology (Academic Press), pp. 48–58.

Figueroa-Clarevega, A., and Bilder, D. (2015). Malignant drosophila tumors interrupt insulin signaling to induce cachexia-like wasting. Developmental Cell 33, 47–55.

Fukuzawa, R., Anaka, M.R., Morison, I.M., and Reeve, A.E. (2017). The developmental programme for genesis of the entire kidney is recapitulated in Wilms tumour. PLoS ONE 12.

Genovese, S., Clement, R., Gaultier, C., Besse, F., Narbonne-Reveau, K., Daian, F., Foppolo, S., Luis, N.M., and Maurange, C. (2019). Coopted temporal patterning governs cellular hierarchy, heterogeneity and metabolism in Drosophila neuroblast tumors. Elife 8.

Glasheen, B.M., Kabra, A.T., and Page-McCaw, A. (2009). Distinct functions for the catalytic and hemopexin domains of a Drosophila matrix metalloproteinase. Proc Natl Acad Sci U S A 106, 2659–2664.

Goveia, J., Rohlenova, K., Taverna, F., Treps, L., Conradi, L.C., Pircher, A., Geldhof, V., de Rooij, L.P.M.H., Kalucka, J., Sokol, L., et al. (2020). An Integrated Gene Expression Landscape Profiling Approach to Identify Lung Tumor Endothelial Cell Heterogeneity and Angiogenic Candidates. Cancer Cell 37, 21-36.e13.

Honegger, B., Galic, M., Kohler, K., Wittwer, F., Brogiolo, W., Hafen, E., and Stocker, H. (2008). Imp-L2, a putative homolog of vertebrate IGF-binding protein 7, counteracts insulin signaling in Drosophila and is essential for starvation resistance. J Biol 7, 10.

Hou, S.X., and Singh, S.R. (2017). Stem-Cell-Based Tumorigenesis in Adult Drosophila. Curr Top Dev Biol 121, 311–337.

Howard, A.M., LaFever, K.S., Fenix, A.M., Scurrah, C.R., Lau, K.S., Burnette, D.T., Bhave, G., Ferrell, N., and Page-McCaw, A. (2019). DSS-induced damage to basement membranes is repaired by matrix replacement and crosslinking. J Cell Sci 132.

Hu, D.J.K., Yun, J., Elstrott, J., and Jasper, H. (2021). Non-canonical Wnt signaling promotes directed migration of intestinal stem cells to sites of injury. Nature Communications 12.

Hwang, N.S., Varghese, S., and Elisseeff, J. (2008). Controlled differentiation of stem cells. In Advanced Drug Delivery Reviews, pp. 199–214.

Jiang, H., Patel, P.H., Kohlmaier, A., Grenley, M.O., McEwen, D.G., and Edgar, B.A. (2009). Cytokine/Jak/Stat Signaling Mediates Regeneration and Homeostasis in the Drosophila Midgut. Cell 137, 1343–1355.

Jiang, H., Tian, A., and Jiang, J. (2016). Intestinal stem cell response to injury: lessons from Drosophila. Cell Mol Life Sci 73, 3337–3349.

Jögi, A., Vaapil, M., Johansson, M., and Påhlman, S. (2012). Cancer cell differentiation heterogeneity and aggressive behavior in solid tumors. In Upsala Journal of Medical Sciences, pp. 217–224.

Kelleher, F.C., Fennelly, D., and Rafferty, M. (2006). Common critical pathways in embryogenesis and cancer. Acta Oncol 45, 375–388.

Kim, J., Chuang, H.C., Wolf, N.K., Nicolai, C.J., Raulet, D.H., Saijo, K., and Bilder, D. (2021). Tumor-induced disruption of the blood-brain barrier promotes host death. Developmental Cell 56, 2712-2721.e2714.

Kim, N., Kim, H.K., Lee, K., Hong, Y., Cho, J.H., Choi, J.W., Lee, J.I., Suh, Y.L., Ku, B.M., Eum, H.H., et al. (2020). Single-cell RNA sequencing demonstrates the molecular and cellular reprogramming of metastatic lung adenocarcinoma. Nature Communications 11.

Korzelius, J., Naumann, S.K., Loza-Coll, M.A., Chan, J.S.K., Dutta, D., Oberheim, J., Gläßer, C., Southall, T.D., Brand, A.H., Jones, D.L., et al. (2014). Escargot maintains stemness and suppresses differentiation in Drosophila intestinal stem cells. The EMBO Journal 33, 2967–2982.

Krakhmal, N.V., Zavyalova, M.V., Denisov, E.V., Vtorushin, S.V., and Perelmuter, V.M. (2015). Cancer Invasion: Patterns and Mechanisms. Acta Naturae 7, 17–28.

Krieger, T., and Simons, B.D. (2015). Dynamic stem cell heterogeneity. In Development (Cambridge) (Company of Biologists Ltd), pp. 1396–1406.

Kwon, Y., Song, W., Droujinine, I.A., Hu, Y., Asara, J.M., and Perrimon, N. (2015). Systemic organ wasting induced by localized expression of the secreted Insulin/IGF antagonist ImpL2. Developmental Cell 33, 36–46.

Lawson, D.A., Kessenbrock, K., Davis, R.T., Pervolarakis, N., and Werb, Z. (2018). Tumour heterogeneity and metastasis at single-cell resolution. Nat Cell Biol 20, 1349–1360.

Lee, J., Cabrera, A.J.H., Nguyen, C.M.T., and Kwon, Y.V. (2020). Dissemination of Ras V12-transformed cells requires the mechanosensitive channel Piezo. Nature Communications 11.

Lee, J., Ng, K.G., Dombek, K.M., Eom, D.S., and Kwon, Y.V. (2021). Tumors overcome the action of the wasting factor ImpL2 by locally elevating Wnt/Wingless. Proc Natl Acad Sci U S A 118.

Lee, S.H., Park, J.S., Kim, Y.S., Chung, H.Y., and Yoo, M.A. (2012). Requirement of matrix metalloproteinase-1 for intestinal homeostasis in the adult Drosophila midgut. Experimental Cell Research 318, 670–681.

Li, H., and Jasper, H. (2016). Gastrointestinal stem cells in health and disease: from flies to humans. Dis Model Mech 9, 487–499.

Liu, Y., Saavedra, P., and Perrimon, N. (2022). Cancer cachexia: lessons from Drosophila. Dis Model Mech 15.

Lodge, W., Zavortink, M., Golenkina, S., Froldi, F., Dark, C., Cheung, S., Parker, B.L., Blazev, R., Bakopoulos, D., Christie, E.L., et al. (2021). Tumor-derived MMPs regulate cachexia in a Drosophila cancer model. Dev Cell 56, 2664–2680 e2666.

Lowell, S., Jones, P., Le Roux, I., Dunne, J., and Watt, F.M. (2000). Stimulation of human epidermal differentiation by delta-notch signalling at the boundaries of stem-cell clusters. Curr Biol 10, 491–500.

Loza-Coll, M.A., Southall, T.D., Sandall, S.L., Brand, A.H., and Jones, D.L. (2014). Regulation of Drosophila intestinal stem cell maintenance and differentiation by the transcription factor Escargot. The EMBO Journal 33, 2983–2996.

Ma, X., Chen, Y., Xu, W., Wu, N., Li, M., Cao, Y., Wu, S., Li, Q., and Xue, L. (2015). Impaired Hippo signaling promotes Rho1-JNK-dependent growth. Proceedings of the National Academy of Sciences of the United States of America 112, 1065–1070.

Markstein, M., Dettorre, S., Cho, J., Neumuller, R.A., Craig-Muller, S., and Perrimon, N. (2014). Systematic screen of chemotherapeutics in Drosophila stem cell tumors. Proc Natl Acad Sci U S A 111, 4530–4535.

Martín-Blanco, E., Gampel, A., Ring, J., Virdee, K., Kirov, N., Tolkovsky, A.M., and Martinez-Arias, A. (1998). puckered encodes a phosphatase that mediates a feedback loop regulating JNK activity during dorsal closure in Drosophila.

Micchelli, C.A., and Perrimon, N. (2006). Evidence that stem cells reside in the adult Drosophila midgut epithelium. Nature 439, 475–479.

Miguel-Aliaga, I., Jasper, H., and Lemaitre, B. (2018). Anatomy and Physiology of the Digestive Tract of Drosophila melanogaster. Genetics 210, 357–396.

Montell, D.J. (2003). Border-cell migration: The race is on. In Nature Reviews Molecular Cell Biology, pp. 13–24.

Oh, H., and Irvine, K.D. (2009). In vivo analysis of Yorkie phosphorylation sites. Oncogene 28, 1916–1927.

Ohlstein, B., and Spradling, A. (2006). The adult Drosophila posterior midgut is maintained by pluripotent stem cells. Nature 439, 470–474.

Pagliarini, R.A., and Xu, T. (1991). A Genetic Screen in Drosophila for Metastatic Behavior. In Proc Natl Acad Sci USA (Weissman), pp. 143–143.

Pan, D. (2010). The hippo signaling pathway in development and cancer. Dev Cell 19, 491–505.

Pandya, P., Orgaz, J.L., and Sanz-Moreno, V. (2017). Modes of invasion during tumour dissemination. Mol Oncol 11, 5–27.

Parent, J.M., Valentin, V.V., and Lowenstein, D.H. (2002). Prolonged seizures increase proliferating neuroblasts in the adult rat subventricular zone-olfactory bulb pathway. J Neurosci 22, 3174–3188.

Peercy, B.E., and Starz-Gaiano, M. (2020). Clustered cell migration: Modeling the model system of Drosophila border cells. In Seminars in Cell and Developmental Biology (Elsevier Ltd), pp. 167–176.

Petruzzelli, M., and Wagner, E.F. (2016). Mechanisms of metabolic dysfunction in cancer-associated cachexia. Genes Dev 30, 489–501.

Porporato, P.E. (2016). Understanding cachexia as a cancer metabolism syndrome. Oncogenesis 5, e200.

Rangarajan, A., Talora, C., Okuyama, R., Nicolas, M., Mammucari, C., Oh, H., Aster, J.C., Krishna, S., Metzger, D., Chambon, P., et al. (2001). Notch signaling is a direct determinant of keratinocyte growth arrest and entry into differentiation. EMBO J 20, 3427–3436.

Song, W., Kir, S., Hong, S., Hu, Y., Wang, X., Binari, R., Tang, H.W., Chung, V., Banks, A.S., Spiegelman, B., et al. (2019). Tumor-Derived Ligands Trigger Tumor Growth and Host Wasting via Differential MEK Activation. Developmental Cell 48, 277-286.e276.

Stevens, L.J., and Page-McCaw, A. (2012). A secreted MMP is required for reepithelialization during wound healing. Mol Biol Cell 23, 1068–1079.

Tu, B., Yao, J., Ferri-Borgogno, S., Zhao, J., Chen, S., Wang, Q., Yan, L., Zhou, X., Zhu, C., Bang, S., et al. (2019). YAP1 oncogene is a context-specific driver for pancreatic ductal adenocarcinoma. JCI Insight 4.

Uhlirova, M., and Bohmann, D. (2006). JNK- and Fos-regulated Mmp1 expression cooperates with Ras to induce invasive tumors in Drosophila. EMBO J 25, 5294–5304.

Vasilyev, A., Liu, Y., Mudumana, S., Mangos, S., Lam, P.Y., Majumdar, A., Zhao, J., Poon, K.L., Kondrychyn, I., Korzh, V., et al. (2009). Collective cell migration drives morphogenesis of the kidney nephron. PLoS Biology 7.

Wang, C., Yin, M.X., Wu, W., Dong, L., Wang, S., Lu, Y., Xu, J., Wu, W., Li, S., Zhao, Y., et al. (2016). Taiman acts as a coactivator of Yorkie in the Hippo pathway to promote tissue growth and intestinal regeneration. Cell Discovery 2.

Weng, A.P., Millholland, J.M., Yashiro-Ohtani, Y., Arcangeli, M.L., Lau, A., Wai, C., Del Bianco, C., Rodriguez, C.G., Sai, H., Tobias, J., et al. (2006). c-Myc is an important direct target of Notch1 in T-cell acute lymphoblastic leukemia/lymphoma. Genes Dev 20, 2096–2109.

Wittkorn, E., Sarkar, A., Garcia, K., Kango-Singh, M., and Singh, A. (2015). The Hippo pathway effector Yki downregulates Wg signaling to promote retinal differentiation in the Drosophila eye. Development (Cambridge) 142, 2002–2013.

Wu, F., Fan, J., He, Y., Xiong, A., Yu, J., Li, Y., Zhang, Y., Zhao, W., Zhou, F., Li, W., et al. (2021). Single-cell profiling of tumor heterogeneity and the microenvironment in advanced non-small cell lung cancer. Nature Communications 12.

Yang, S.A., Portilla, J.M., Mihailovic, S., Huang, Y.C., and Deng, W.M. (2019). Oncogenic Notch Triggers Neoplastic Tumorigenesis in a Transition-Zone-like Tissue Microenvironment. Dev Cell 49, 461–472 e465.

Yeo, S.K., Zhu, X., Okamoto, T., Hao, M., Wang, C., Lu, P., Lu, L.J., and Guan, J.L. (2020). Single-cell RNA-sequencing reveals distinct patterns of cell state heterogeneity in mouse models of breast cancer. eLife 9, 1–24.

Yeom, E., Shin, H., Yoo, W., Jun, E., Kim, S., Hong, S.H., Kwon, D.W., Ryu, T.H., Suh, J.M., Kim, S.C., et al. (2021). Tumour-derived Dilp8/INSL3 induces cancer anorexia by regulating feeding neuropeptides via Lgr3/8 in the brain. Nat Cell Biol 23, 172–183.

Zakrzewski, W., Dobrzynski, M., Szymonowicz, M., and Rybak, Z. (2019). Stem cells: Past, present, and future. In Stem Cell Research and Therapy (BioMed Central Ltd.).

Zeng, X., and Hou, S.X. (2015). Enteroendocrine cells are generated from stem cells through a distinct progenitor in the adult Drosophila posterior midgut. Development 142, 644–653.

Zhai, Z., Kondo, S., Ha, N., Boquete, J.P., Brunner, M., Ueda, R., and Lemaitre, B. (2015). Accumulation of differentiating intestinal stem cell progenies drives tumorigenesis. Nature Communications 6.

