## Supplemental figures (Revision) for "The native cell differentiation program aberrantly recapitulated in *yki*^*3S*/*A*^-induced intestinal hyperplasia drives invasiveness and cachexia-like wasting phenotypes"

**Supplemental Information (Pranoto et al.)**

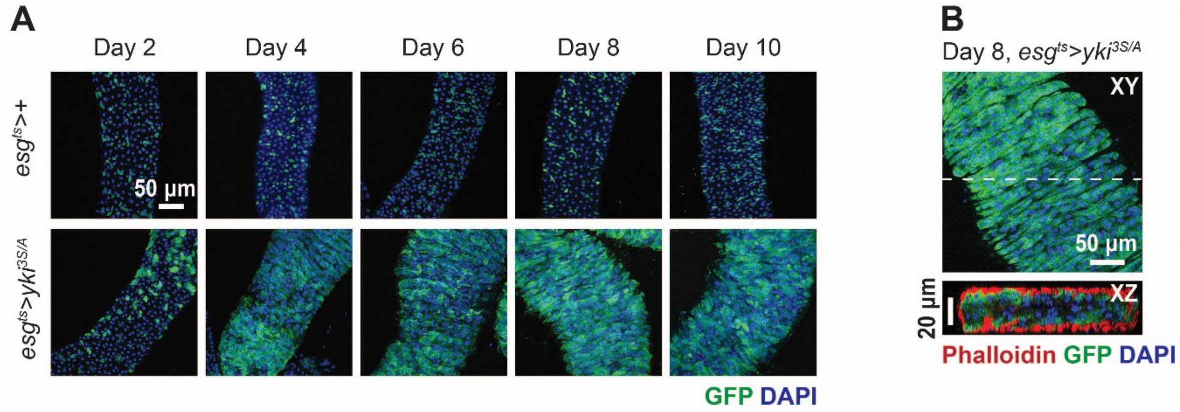


**Figure S1. Expression of *yki*$^{3S/A}$ in ISCs and EBs induces hyperplasia in the midgut.**

**A.** Posterior region of *esg*$^{ts}$>+ and *esg*$^{ts}$ >*yki*$^{3S/A}$ midguts. *yki*$^{3S/A}$ was expressed with *esg*$^{ts}$ by shifting to 29°C for the indicated durations. The cells manipulated by *esg*$^{ts}$ are marked with GFP (green), and nuclei are stained with DAPI (blue). **B.** Representative orthogonal views of the posterior midguts at day 8 of induction.

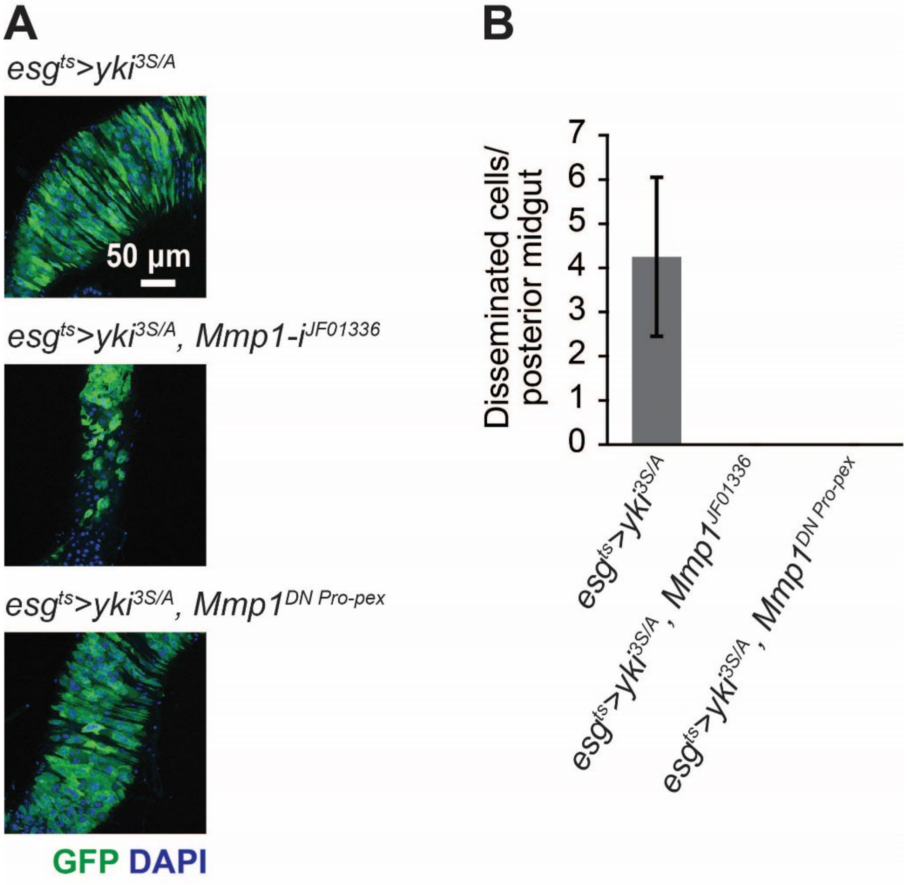


**Figure S2. Mmp1 knockdown or inhibition suppresses the growth of *yki*$^{3S/A}$ midgut tumors and dissemination of *yki*$^{3S/A}$ cells.**

**A.** Posterior midguts after inducing transgenes for 12 days. *JF01336* was used to knockdown Mmp1 in *yki*$^{3S/A}$ cells. *Mmp1*$^{DN\ Pro\text{-}pex}$ is a dominant negative *Mmp1* allele. **B.** Quantification of disseminated cells detected on the surface of posterior midgut.

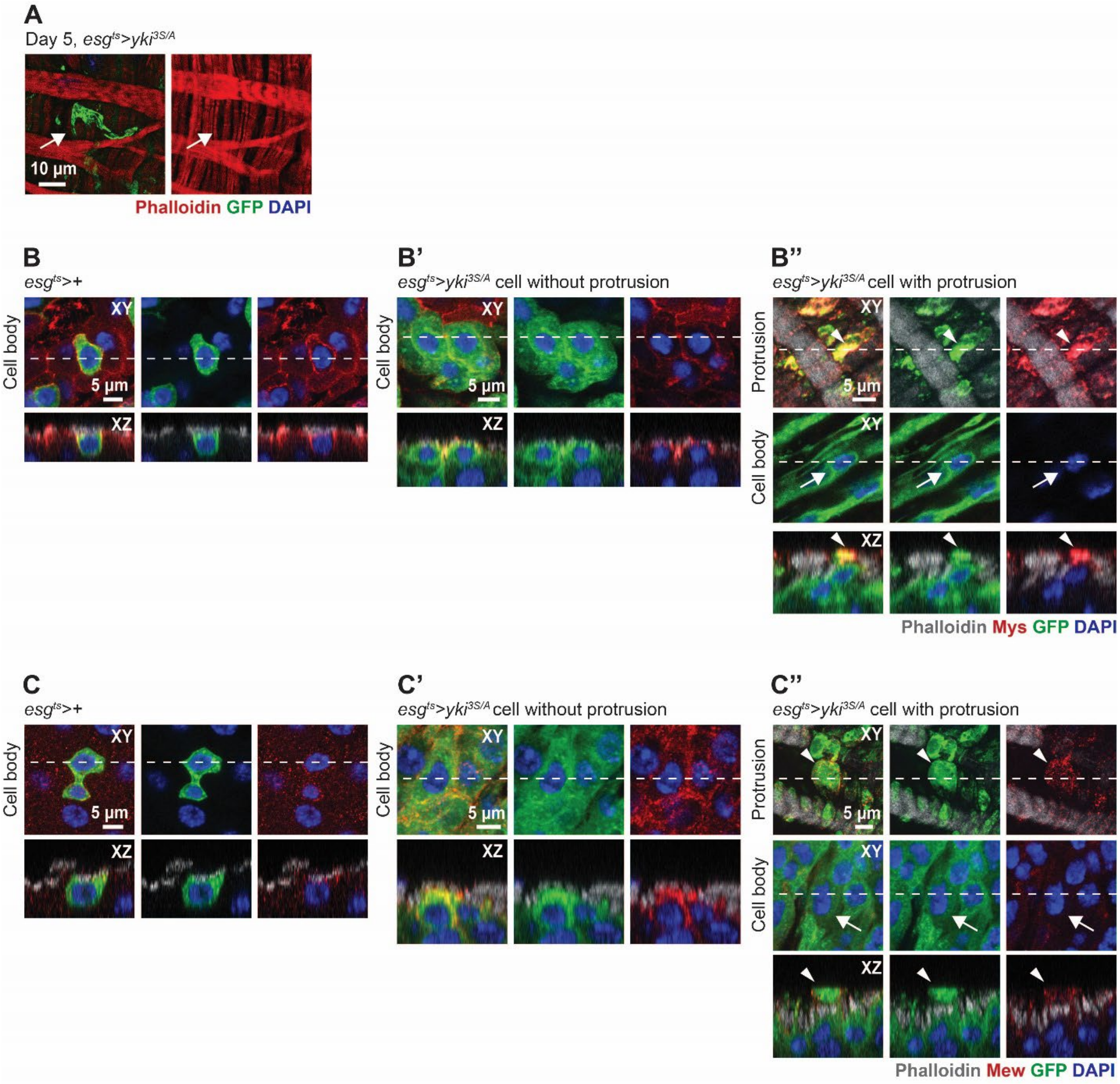


**Figure S3. Focal adhesion components are upregulated in *yki$^{3S/A}$* cells and accumulated at the protruding leading edges.**

**A.** Representative image of a protrusion generated from *yki$^{3S/A}$* cell. Phalloidin signals show the visceral muscle. *yki$^{3S/A}$* cells generate protrusions across the visceral muscle without damaging the tissue. **B-B''.** Cellular localization of Mys (red) in representative *esg$^{ts}$* cell (**B**), *yki$^{3S/A}$* cell without protrusion (**B'**), and *yki$^{3S/A}$* cell with protrusion (**B''**).

Arrowhead is used to indicate protrusion observed at the surface of the visceral muscle. Arrow is used to indicate the body of *yki*$^{3S/A}$ cell generating protrusion. **C-C''.** Cellular localization of Mew (red) in representative *esg*$^{ts}$ cell (**C**), *yki*$^{3S/A}$ cell without protrusion (**C'**), and *yki*$^{3S/A}$ cell with protrusion (**C''**). Arrowhead is used to indicate protrusion observed at the surface of the visceral muscle. Arrow is used to indicate the body of *yki*$^{3S/A}$ cell generating protrusion.

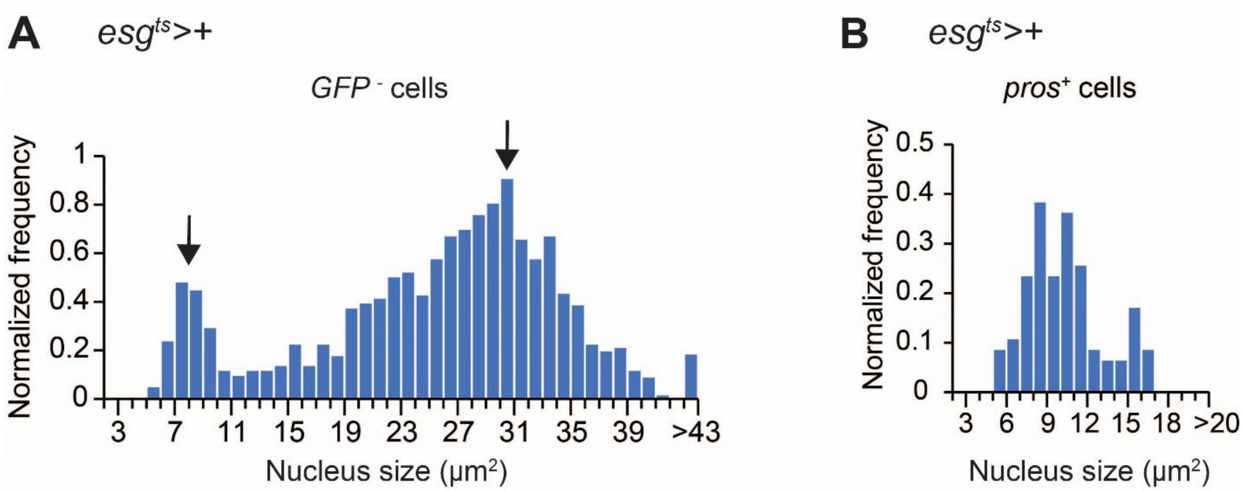


**Figure S4. Nuclear size distribution of EEs and ECs in the posterior midguts.**

**A.** Nuclear size distribution of *GFP*$^{-}$ cells (ECs and EEs) in the posterior midguts at day 6. GFP is driven by *esg*$^{ts}$. Arrows indicate two peaks. N=2013. **B.** Nucleus size distribution of *pros*$^{+}$ cells at day 6. N=100.

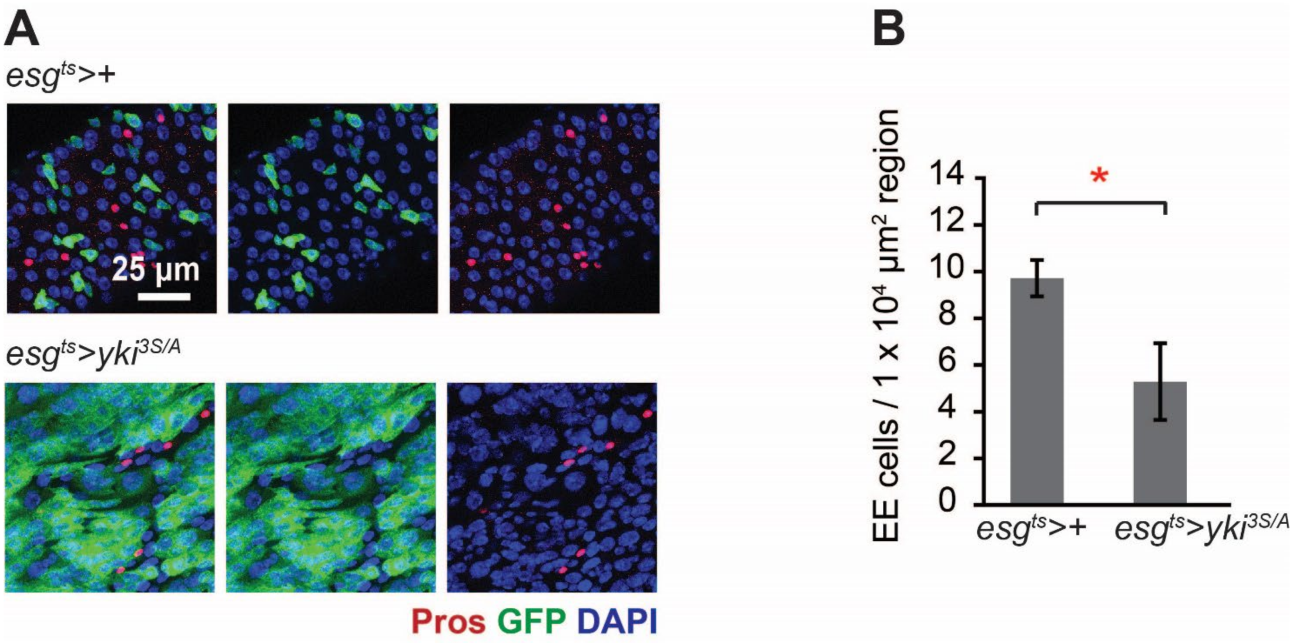


**Figure S5. Expression of *yki*$^{3S/A}$ with *esg*$^{ts}$ accumulates cells in EC lineage in the expense of EE lineage.**

**A.** Representative images of Pros staining (red) in the upper region of the posterior midguts. Transgenes were induced for 8 days. Cells manipulated by *esg*$^{ts}$ are marked with GFP (green). **B.** Quantification of *pros*$^{+}$ cells in the upper region of the posterior midguts. *yki*$^{3S/A}$-induced hyperplasia are more prominent in the upper region of the posterior midgut. Thus, we sampled from the upper region for quantification. N=7 midguts for each genotype. Mean±SEMs are shown. *P<0.05, two-tailed unpaired Student's t-test.

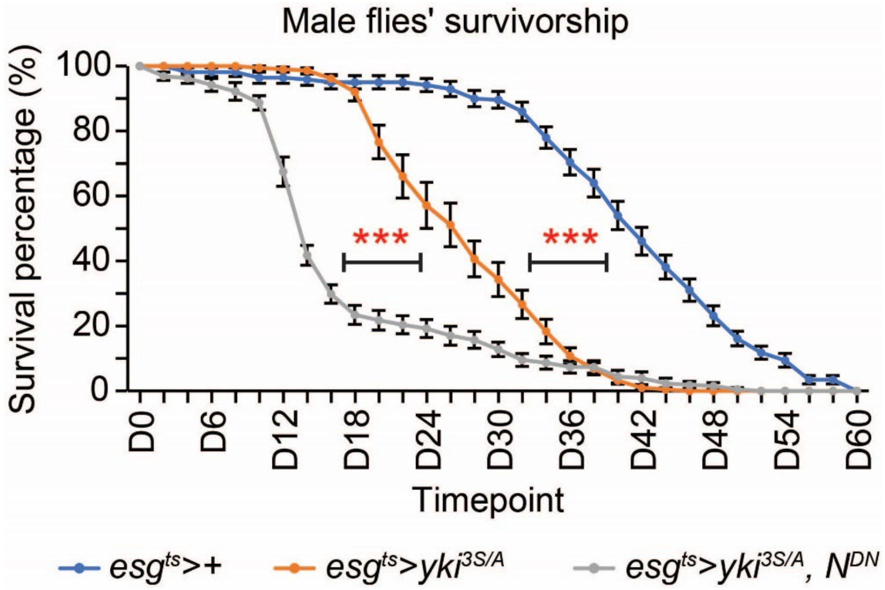


**Figure S6. Blocking the EC differentiation in *yki*$^{3S/A}$ midgut tumors reduces the lifespan of tumor-bearing male flies.**

**A.** Survivorship in male flies. *esg*$^{ts}$>+ intestine, blue, N=198 flies in 20 replicates; *esg*$^{ts}$>*yki*$^{3S/A}$, orange, N=203 flies in 18 replicates; *esg*$^{ts}$>*yki*$^{3S/A}$, *N*$^{DN}$, gray, N=210 flies in 20 replicates. Mean±SEMs are shown. ***p < 0.001, log-rank test.

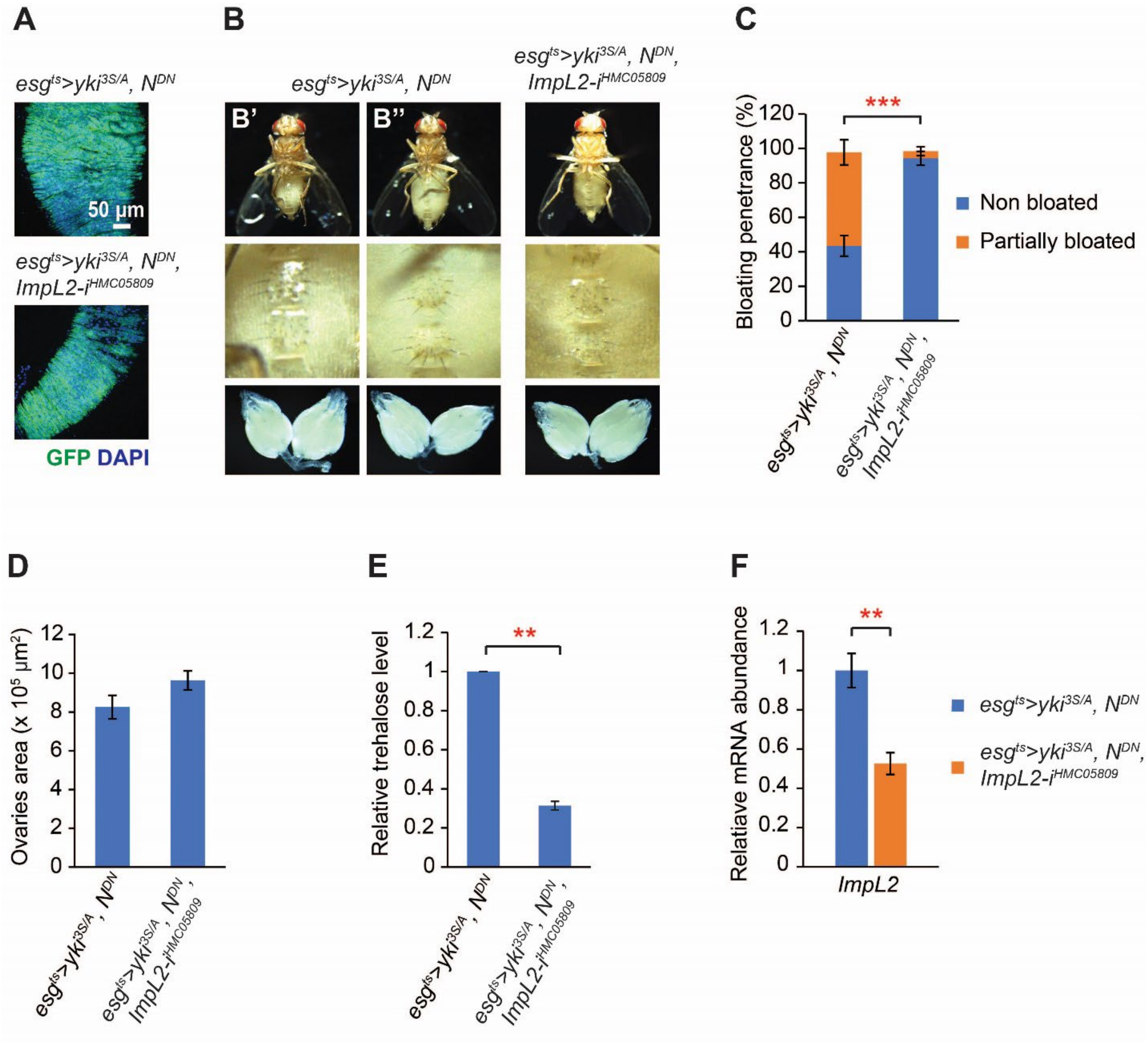


**Figure S7. Depletion of *ImpL2* in *yki*$^{3S/A}$, *N*$^{DN}$ tumors further suppresses the wasting phenotypes.**

**A.** Representative images of *esg*$^{ts}$*>yki*$^{3S/A}$*, N*$^{DN}$ and *esg*$^{ts}$*>yki*$^{3S/A}$*, N*$^{DN}$, *ImpL2-i*$^{HMC05809}$ midgut tumors. Transgenes were induced for 8 days. Tumor cells are marked by GFP (green). **B.** Representative ventral views of flies and ovary images. Upper panels, ventral views; middle panels, magnified views of the corresponding fly's abdominal area; lower panels, images of the corresponding fly's ovaries. The upper panels in

*$esg^{ts}$>$yki^{3S/A}$*, *$N^{DN}$* show representative non-bloated (normal state) (**B'**) and partially bloated (**B''**) fly's abdominal views. Transgenes were induced for 8 days. **C.** Quantification of bloating syndrome penetrance. Transgenes were induced for 8 days. N=272 animals (*$esg^{ts}$>$yki^{3S/A}$, $N^{DN}$*) and N=182 animals (*$esg^{ts}$>$yki^{3S/A}$, $N^{DN}$*, *ImpL2-$i^{HMC05809}$*). Mean±SEMs are shown. ***P<0.001, chi-square test. **D.** Quantification of ovaries size. N=20 pairs of ovaries) (*$esg^{ts}$>$yki^{3S/A}$, $N^{DN}$*) and N=21 pairs of ovaries (*$esg^{ts}$>$yki^{3S/A}$, $N^{DN}$*, *ImpL2-$i^{HMC05809}$*). Mean±SEMs are shown. Two-tailed unpaired Student's t-test. **E.** Relative trehalose levels in *$esg^{ts}$>$yki^{3S/A}$, $N^{DN}$* and *$esg^{ts}$>$yki^{3S/A}$, $N^{DN}$, ImpL2-$i^{HMC05809}$* flies at day 8 of transgene induction. Mean±SEMs are shown. N=6 female flies, 3 biological replicates. **P<0.01, two-tailed unpaired Student's t-test. **F.** Relative *ImpL2* mRNA levels. Depletion of ImpL2 further reduced *ImpL2* mRNA levels in *$yki^{3S/A}$, $N^{DN}$*, *ImpL2-$i^{HMC05809}$* tumors compared to *$yki^{3S/A}$, $N^{DN}$* tumors. N=10 midguts, 3 biological replicates. **P<0.01, two-tailed unpaired Student's t-test.

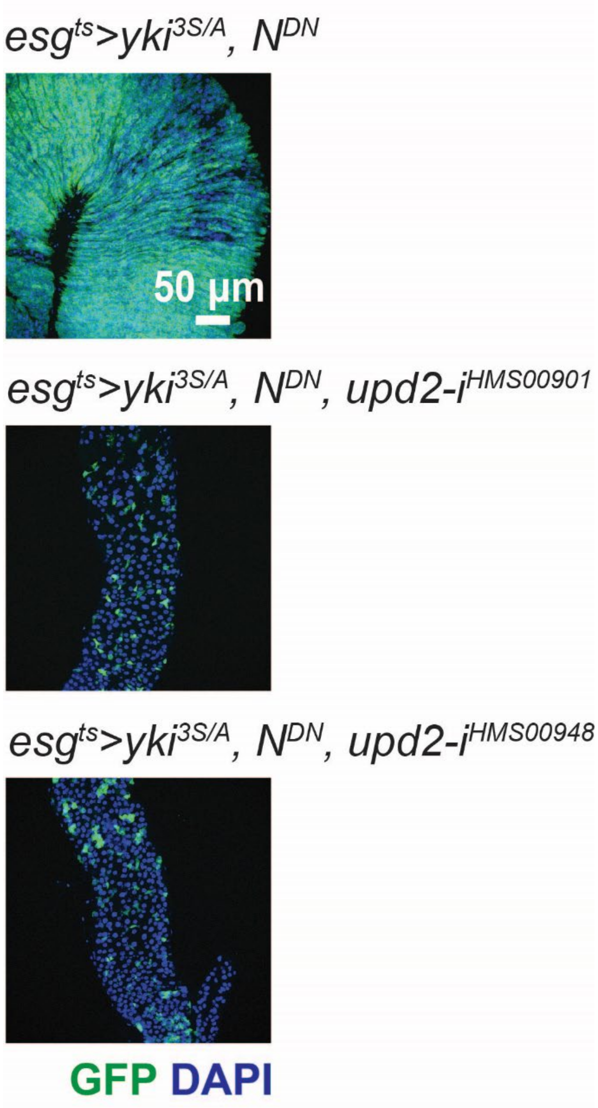


**Figure S8. Upd2 expressed in *yki*$^{3S/A}$, *N*$^{DN}$ cells is required for the tumor development.**

Representative images of the posterior midguts are shown. Two independent *upd2* RNAi lines (*HMS00901* and *HMS00948*) are used to knockdown *upd2* in *yki*$^{3S/A}$, *N*$^{DN}$ cells for 8 days. Requirement of Upd2 in *yki*$^{3S/A}$, *N*$^{DN}$ cells for the tumor development indicates that Upd2 is expressed in the tumor cells.
